# The Last Iberian Record of Eurasian Lynx (*Lynx lynx*): Osteometry and Historical Implications of the Lynx from Sima Topinoria (Cantabria, Spain)

**DOI:** 10.64898/2025.12.15.694219

**Authors:** Eva Fernández-Bejarano, Carlos Nores Quesada, Fernando Serrulla Rech, Borja Palacios Alberti, Juan Martín Otero, Gloria González-Fortes, Aurora Grandal-d’Anglade

## Abstract

The Eurasian lynx (*Lynx lynx*) was historically native to the Iberian Peninsula, as evidenced by scarce paleontological records and sightings across northern Spain, dating from the Last Glacial Maximum until the 17th century. A novel and nearly complete skeleton of a medium-sized felid, morphologically identified as *L. lynx*, was recently recovered from Sima Topinoria in Picos de Europa (Cantabria, Spain). The present study aims to recover and analyze the full skeletal assemblage, establishing its chronological framework, taxonomic identification, and comparative craniometry with other ancient and modern Eurasian lynxes. Radiocarbon dating estimates the specimen around 210 years BP, representing the most recent confirmed occurrence of Eurasian lynx in the Iberian Peninsula. This evidence revises the accepted timeline for the species’ extirpation in the region, indicating its persistence into the early 19th century, connecting physical evidence with historical and traditional narratives. Morphometric analysis identifies the individual as an adult male with an estimated body mass of 19.7 kg. Comparative osteometric analyses revealed that the specimen from Sima Topinoria falls within the average size for modern males from the Carpathian population, while being significantly smaller than older Holocene males from the Iberian Peninsula (∼2570 yBP), suggesting a possible trend toward a protohistoric body size reduction in the Holocene, akin to patterns reported for other mammalian fauna from the Cantabrian Mountains. These results redefine the Eurasian lynx’ s historical range collapse in southwestern Europe, suggesting that the species survived until recent times, coinciding with periods of intense anthropogenic landscape change. The study highlights the critical role of paleontological data in refining extinction chronologies and contributes valuable insights into the biogeographic history of this elusive feline in western Europe.

## 1. INTRODUCTION

### 1.1. The Palearctic lynxes

Two lynx species inhabit Eurasia. The Eurasian lynx (*Lynx lynx*) is the largest of the four extant lynx species and one of the most widely distributed felids. Its range extends from Western Europe across Siberia to the Himalayas and the Middle East (von Arx et al., 2004). This large-bodied species is adapted to temperate and boreal forest ecosystems. Historically widespread, the Eurasian lynx experienced a significant contraction of its range during the 19th and 20th centuries due to habitat loss, prey depletion, and direct persecution. By the early 20th century, its westernmost presence had become restricted to regions such as Fennoscandia, the Balkans, and the Carpathians (Linnell et al., 2009; Schmidt et al., 2011).

In contrast, the Iberian lynx (*Lynx pardinus*) is smaller, more specialized, and currently survives as a relict species in the south-westernmost part of Europe, limited to southern areas of the Iberian Peninsula. However, this represents only a fraction of its historical range. Holocene archaeological sites in northern Iberia, including the Cantabrian Mountains and the Pyrenees, have yielded remains attributed to the Iberian lynx (Villaluenga, 2016). During the Pleistocene, a larger form of Iberian lynx, sometimes referred to as *Lynx pardina spelaea* (Boule), has been documented in sites in southern France (Kurtén & Granqvist, 1985; Fosse et al., 2021).

For decades, it was widely believed that the ranges of the Eurasian and Iberian lynx did not overlap in the Iberian Peninsula, where only the latter was thought to have occurred (Kratochvil, 1968; Breitenmoser et al., 2000). Nevertheless, some Holocene sites in the Cantabrian region have yielded lynx remains identified as *Lynx spelaea*, a taxon of uncertain affinity but larger than the modern Iberian lynx (Altuna, 1972; Castaños, 1984).

Recent genetic studies have challenged this assumption. Analyses of Holocene remains from northern Iberia have confirmed the presence of *Lynx lynx* (Rodríguez-Varela et al., 2016), while *Lynx pardinus* has been unexpectedly identified in archaeological sites from Italy (Rodríguez-Varela et al., 2015; Mecozzi et al., 2021). Due to this unexpected and extensive distribution of the Iberian lynx, some authors even suggest changing its vernacular name to ‘Mediterranean lynx’ (Tura-Poch et al., 2022). These findings strongly suggest that, at least during certain Holocene phases, the two species may have coexisted in partially overlapping or adjacent ranges. This scenario has important implications for understanding past interspecific interactions, such as ecological competition, habitat segregation, and the reliability of species identification in fragmented fossil assemblages.

### 1.2. The cerval wolf

Popular tradition in the north of the Iberian Peninsula tells us of a prey animal capable of attacking livestock, similar to a big carnivore with mottled fur, which was called “Lobo cerval*”*, or cerval-wolf (latinized as *Lupus cervarius*), a name that highlights its preference for hunting deer, although historical sources highlight their predation on smaller livestock, such as sheep and goats, even calves (Nores et al., 2015). This vernacular term contrasts with the true wolf (*Canis lupus*), which is more often associated with predation on livestock such as cattle, for which sometimes received the name of “lobo vaquero” or cow-wolf (Nores & Vázquez, 1984). The name ‘cerval wolf’ reflects the lynx’s role as a large solitary predator in forested habitats, sharing some ecological traits with the wolf, but targeting different prey. Mentions of this enigmatic animal are very numerous and can be found from Galicia, in the most north-western part of the peninsula, and all along the Cantabrian Mountain range, reaching the Basque Country and Navarre. These are not old legends, but direct encounters with the animal, or sightings reported by shepherds or hunters from one or two generations ago at most.References to the cerval wolf are even recorded in newspapers, books with local geographical descriptions, and certificates of rewards paid for its death (Nores et al., 2015). The description of the animal and the fact that it was capable of killing large prey, suggested that the cerval wolf could have been a Eurasian lynx (Villalpando-Moreno, 2020).

From the historical citations collected between the 16th and 19th centuries attributable to lynxes in the Iberian Peninsula, Clavero and Delibes (2013) concluded that those included in the Atlantic climate regions, from Galicia to the border of the Pyrenees, where the name ‘cerval wolf’ dominated, would be attributable to the Eurasian lynx *Lynx lynx*, while those collected in the Mediterranean climate, preferably mentioned as a cat, ‘gato cerval’ or ‘gato clavo’, would be better suited to the smaller Iberian lynx *Lynx pardinus*. This interpretation seems to be reinforced by the confirmation of the presence of Eurasian lynx in the Atlantic rim through genetic analysis of fossils (Rodríguez-Varela et al., 2016). However, no physical evidence of the existence of the Eurasian lynx had been found in the Cantabrian Mountain range in the present or in the recent past. The most recent evidence of the species consists of scant remains discovered in the Pyrenees (Sima Serpenteko, Navarre), dated to 412 years BP (Rodríguez-Varela et al., 2016). Outside of the Pyrenees, the most recent among the records is an incomplete skeleton from the Cueva de los Cinchos, in Asturias, dated to roughly 1,750 years BP (Rodríguez-Varela et al., 2016), too ancient for its memory to endure in popular narrative to our days.

### 1.3. The Discovery in Torca de la Topinoria

Located in the heart of the Cantabrian Mountains in northern Spain, the Picos de Europa National Park is one of the most iconic protected areas on the Iberian Peninsula (Fig. 1). It spans 67,455 hectares across the provinces of Asturias, León, and Cantabria, encompassing eleven municipalities. The park’s landscape is dominated by three massive limestone ranges—Western, Central, and Eastern—deeply sculpted by the valleys of the Sella, Cares, and Deva rivers. A wide range of altitudes, combined with centuries of traditional land use, has shaped a remarkably diverse environment that includes vast forests, alpine pastures, and steep rocky crags. This diversity sustains a rich biodiversity, including some of the most emblematic and endangered species of the Cantabrian range. The striking geological features of the park are largely the result of extensive karst processes, which have given rise to numerous caves and chasms—among them, Torca de la Topinoria, the site of the paleontological discovery described below.

**Figure 1.**
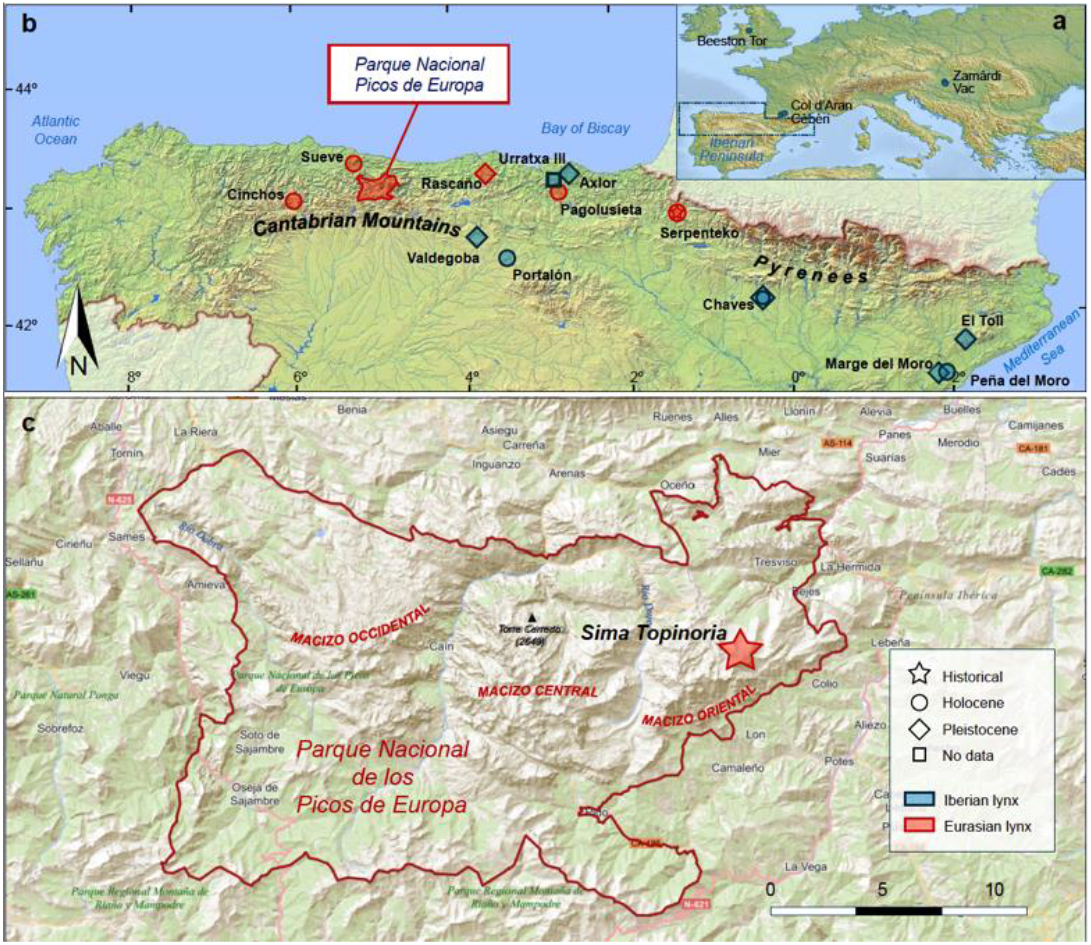
Location of the site Sima Topinoria and other sites with *Lynx* remains mentioned in the text.

Torca de la Topinoria (SN-2), also called Sima Topinoria, is a cave located on the northern slopes of Pico Samelar, within the municipality of Cillorigo de Liébana, in the Eastern Massif of the Picos de Europa (Fig. 1). (Coordinates: X 363063; Y 4786651; Altitude above sea level: 1636 m). The cave was first explored in the 1990s, and in recent years, new exploration campaigns have been carried out to improve its documentation and complete the topographic survey.

In 2020, during one of these re-exploration campaigns, speleologists from Flash Caving Club (Madrid) and Tracalet (Valencia) discovered a nearly complete lynx skeleton that had never been reported before. The animal was found in a side chamber approximately 100 meters deep (see cave map, Fig. 2).

**Figure 2.**
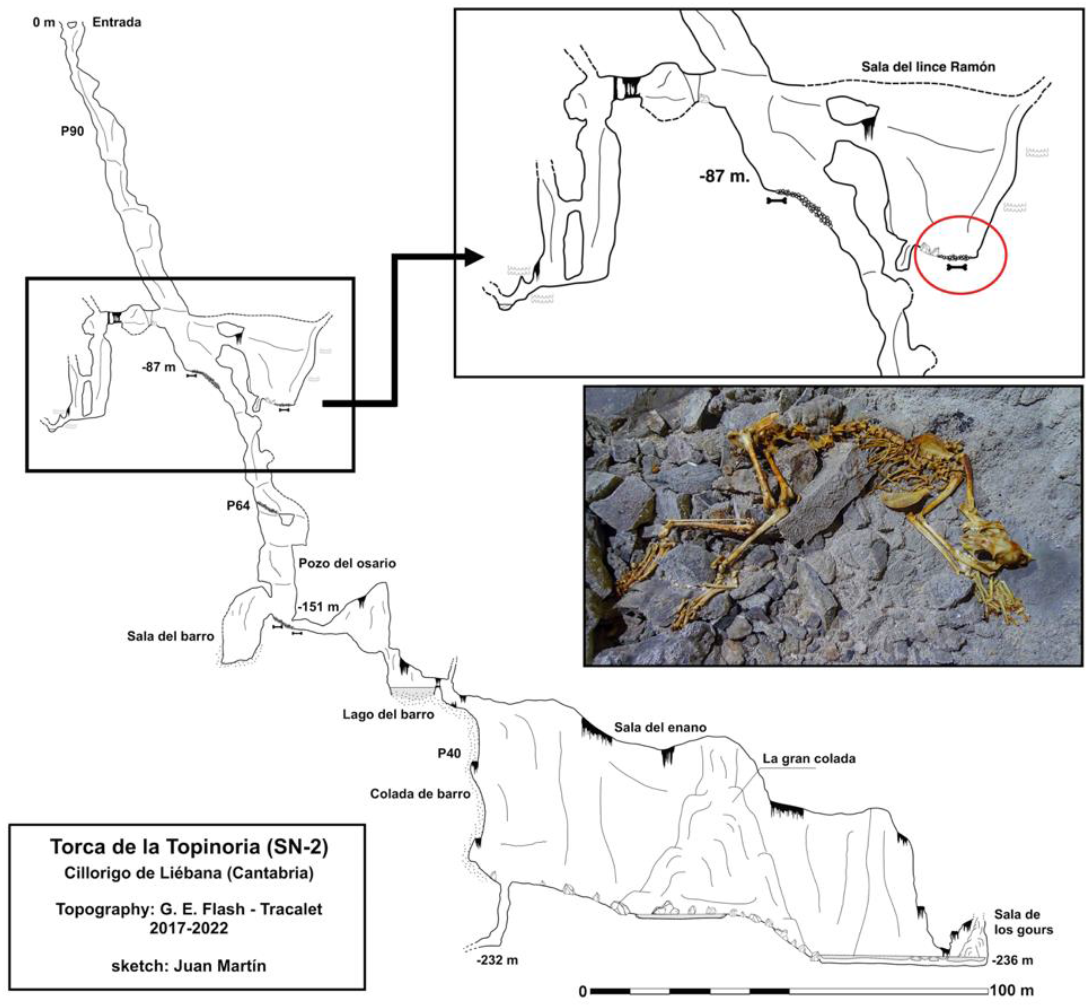
Section of the cave and position of the Lynx skeleton.

The animal was lying on its left side in a natural resting position (See Fig. 2). All skeletal elements were present and articulated, except for the skull and mandibles, which had been slightly displaced by a few decimeters. The canines were missing, likely removed by a previous visitor to the cave. The bones were in excellent condition, showing only minimal oxide staining, suggesting a relatively recent date. After an extensive photographic record was made and several specialists were consulted, the remains were preliminarily identified as those of a Eurasian lynx (*Lynx lynx*). However, a detailed morphological and metric analysis was necessary to confirm the taxonomic identification. Thanks to the generous collaboration of speleologists from Caving Club Flash (Madrid) and the funding provided by the Picos de Europa Natural Park, the skeleton was fully recovered, dated, and subjected to a comprehensive study.

### 1.4. Objectives

In this article, we present a detailed anatomical and metric study of the newly discovered Eurasian lynx skeleton from the Cantabrian Mountains. Our objectives are threefold: (i) to confirm the identification of the specimen as *Lynx lynx* based on specific cranial and postcranial features; (ii) to provide a complete set of osteometric data that can serve as a reference for comparison in future zooarchaeological work; and (iii) to contextualize this finding within the broader framework of Holocene lynx distribution in the Iberian Peninsula, including its implications for conservation biology, biogeography, and historical ecology.

## 2. MATERIAL AND METHODS

### 2.1. Palaeontological data

Given the difficulty of accessing the skeletal remains, a strategy was devised that involved a preliminary study of the position of each bone based on photographs taken by members of the Flash Caving Group (Madrid), who were also responsible for the recovery of the remains. Once inside the sinkhole, the cavers proceeded to carefully extract the bones in an orderly manner, grouped by anatomical sections. This approach allowed for the subsequent reconstruction of the skeleton, with correct assignment of smaller appendicular elements (phalanges) to the appropriate limbs.

At the laboratory of the University of A Coruña (UDC), the bones were cleaned sequentially using warm water to remove the minimal amount of sediment adhered to them. They were then left to dry at a cold and controlled temperature to prevent sudden changes in volume that might cause cracking or fracturing. The bones were in a very good state of preservation, with an almost fresh appearance. Only a few ribs on the left side showed deterioration in the proximal area, which was later identified as pathological. After stabilizing the remains, the bones were identified and organized for labeling and metric analysis (Fig. 3). Anatomical identification was based on the *Osteological Atlas* by Pales and García (1981) and the *Digital Atlas of the Iberian Lynx Skeleton* (https://thevirtualmuseumoflife.com/lynx/index-es.php).

**Figure 3.**
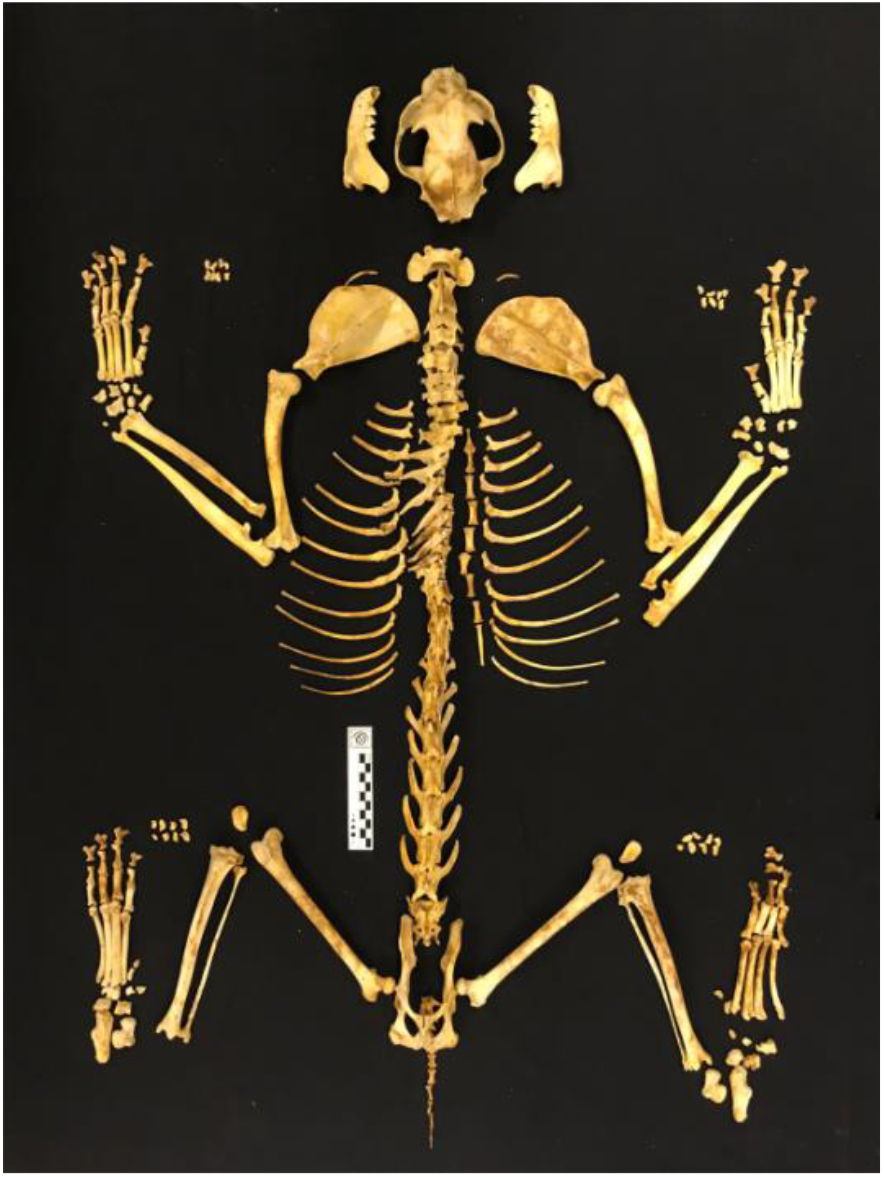
The Eurasian lynx skeleton from Sima Topinoria after reconstruction in the laboratory.

**Figure 4.**
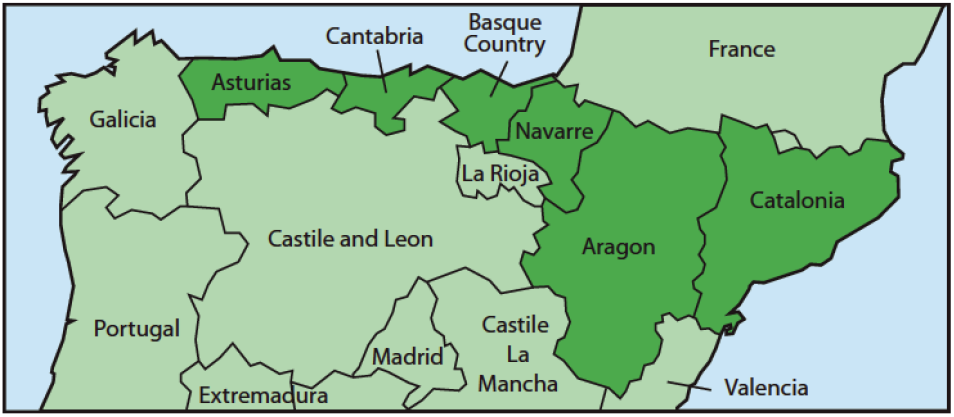
Regions from which historical data on the presence of “lobo cerval” (presumably *Lynx lynx*) were analized.

For radiocarbon dating, the right 11th rib was selected. A 250 mg fragment was extracted and sent to Beta Analytic Inc. laboratory (Miami, USA) where it underwent a classic organic matter extraction treatment (acid-alkali-acid) prior to graphitization and dating by Accelerator Mass Spectrometry (AMS).

Metric data collection followed the standard protocol of Von den Driesch (1976), with additional measurements adapted for felids based on the osteometric system for ursids proposed by Tsoukala and Grandal (2002). The Supplementary Material 1 includes diagrams illustrating all measurements taken. A set of Mitutoyo digital calipers and an osteometric board were used for measuring long bones. Body mass (BM) was estimated from the condylobasal length following published allometric equations for felids (Van Valkenburgh, 1990). The height at the withers was calculated according to Gal et al. (2024). The biological age of the specimen was determined following García Perea et al (1996), based on the sutures between cranial bones and the sutures of long bones in general.

To assess cranial morphological variation among Eurasian lynx populations, total length and zygomatic breadth were analyzed, and a biplot was generated in R v.4.4.3. Craniometric data were compiled for multiple populations of Eurasian lynx, including a few prehistoric specimens previously published in the literature (Boule & Villenueve, 1927; Ognev, 1962; Stollmann, 1963; Vasiliu & Decei, 1964; de Beaufort, 1965; Stroganov, 1969; Clot & Besson, 1974; Altuna, 1980; Heptner & Sludskij, 1980; Miric, 1981; Andersen & Wiig, 1986; Bartosiewiez, 1993; Červený & Koubek, 2000; Gomercic et al., 2010; Albayrak, 2012; Dayan et al., 2017; Mecozzi et al., 2021; Bizhanova & Grachev, 2022; Gal et al., 2022). We also include some metric data from another Eurasian lynx skull from Sierra del Sueve (Asturias, Spain) dated to 4705 yBP that had only been partially published (Nores, 1999).

For the study of the pathological ribs, a morphoscopic study and conventional radiography were performed using a Siemens Multixtop digital device at 48 kV and 4 mA. The images were obtained using the Rain Alma viewer and downloaded in jpeg format (Herring, 2020).

Following its metric study, the lynx skeleton was deposited at the Museum of Prehistory and Archaeology of Cantabria (MUPAC), with registration number 2297/2023 and exhibition number F-4025.

### 2.2. Historical data

Information on changes in historical distribution comes from different geographical dictionaries in which territorial descriptions often also provide lists of fauna (game or vermin) (see Nores & López Bao, 2022). The period we have considered is from the beginning of the 19th century (contemporary to the specimen studied) to the middle of the 20th century.

Data from the beginning of the 19th century are restricted to some complete regions of the Cantabrian Mountains and part of the Pyrenees (4). The Cantabrian regions: Asturias (Martínez Marina 1801-1802) and the Basque Country (Anonymous, 1802), have faunal information from this period in almost all current municipalities, so this information has no apparent geographical bias. The Pyrenean regions have more limited information. Thus, Navarre provides faunal information for only 9 present-day municipalities spread throughout the region (Anonymous, 1802); Aragon only for part of the provinces of Huesca and Zaragoza (Sumán, 1802 [2015]) and Catalonia for part of Lleida, Barcelona and Girona (Zamora, 1788-1790, in Maluquer i Sostres, 1992).

Faunistic information from the mid-19th century and mid-20th century exists for the whole of Spain (Madoz, 1846-1850; Bleiberg & Quirós, 1956-1961), but only the municipalities mentioned in the sources from the beginning of the 19th century were taken into account to avoid biases from other areas with different characteristics that would make comparison difficult.

The information collected was referenced to current municipalities in order to statistically compare changes between municipalities with presence/non-detection of lynx using Fisher’s Exact Probability test with a two-tailed test in a 2x2 (PEF2x2) and 2x3 (PEF2x3) contingency table using the online statistical package VassarStats (http://vassarstats.net/). Global changes were only assessed in a 2x3 contingency table in cases where there could be a generalized trend in frequencies over the three points in time assessed, and no significant changes were detected between two periods that could be attributed to the low number of data available (N < 300), as in the case of Navarre and in some other cases where it might be useful for a more correct interpretation of the results.

## 3. RESULTS

### 3.1. CHRONOLOGY

The AMS ^14^C dating yields a conventional age of 210 ± 30 years BP (Beta–644857). Calibration was performed using a Bayesian statistical program (Bronk Ramsey, 2009), which incorporates all variables, including the dating margin of error (± 30 years). The calibration curve used was IntCal20 (Reimer et al., 2020). As a result, the calibrated age of the lynx remains, with a 95.4% probability, falls within the calendar date ranges shown in Table 1.

**Table 1.** Radiocarbon dating of the *Lynx lynx* from Sima Topinoria, calibrated with the curve IntCal20 (Reimer et al., 2020)

| Radiocarbon dating |  |
| --- | --- |
| Probability | Calibrated calendar age |
| (51.6%) | 1729 – 1808 cal AD |
| (29.5%) | 1642 – 1688 cal AD |
| (14.3%) | 1924 – Post AD 1950 |

Therefore, the remains most likely date to the 18^th^ century (51.6% probability), although there is a smaller probability that they are slightly older (second half of the 17th century) or even subrecent (20^th^ century). Apart from this specimen, the most recent known Eurasian lynx remains in the Iberian Peninsula come from the Serpenteko sinkhole in Navarre (Pyrenees), where a dated bone fragment yielded a calibrated age between 505 and 319 years BP, corresponding to a calendar age between the mid-15^th^ and mid-16^th^ centuries (Rodríguez-Varela et al., 2016).

### 3.2. Taxonomy

The specific identification of the specimen is based on cranial morphological features. Morphologically, the skull displays several diagnostic traits:

- **Shape of the temporal crests. V-shaped temporal crests**, lacking a “lyre” shape, is a characteristic of the Eurasian lynx morphotype (García-Perea, 1996) (Fig. 5a). The sagittal crest is, however, shorter and more gracile than other *L. lynx* specimens found in the Iberian Peninsula, recovered from Pagolusieta, Navarre (2,570 ± 30 yBP)
- **Separation between the foramina on the posterior margin of the tympanic bulla**. In *Lynx pardinus*, both the hypoglossal and jugular foramina converge into a single fossa, while in *Lynx lynx* foramina are clearly separated into distinct cavities. In the case of the Topinoria specimen, foramen morphology is consistent with *L. lynx* (Fig. 5f) (Mecozzi et al., 2021).
- Dentition also exhibits taxonomically informative characteristics, particularly the presence of a small metaconid on the lower carnassial. This feature is common in most populations of L. lynx but occurs only rarely in *L. pardinus* (García-Perea, 1985; Kitchener et al., 2017; Mecozzi et al., 2021). The Topinoria specimen displays a small yet distinct metaconid (Fig. 6d and 6f), further supporting its identification as *Lynx lynx*.

**Figure 5.**
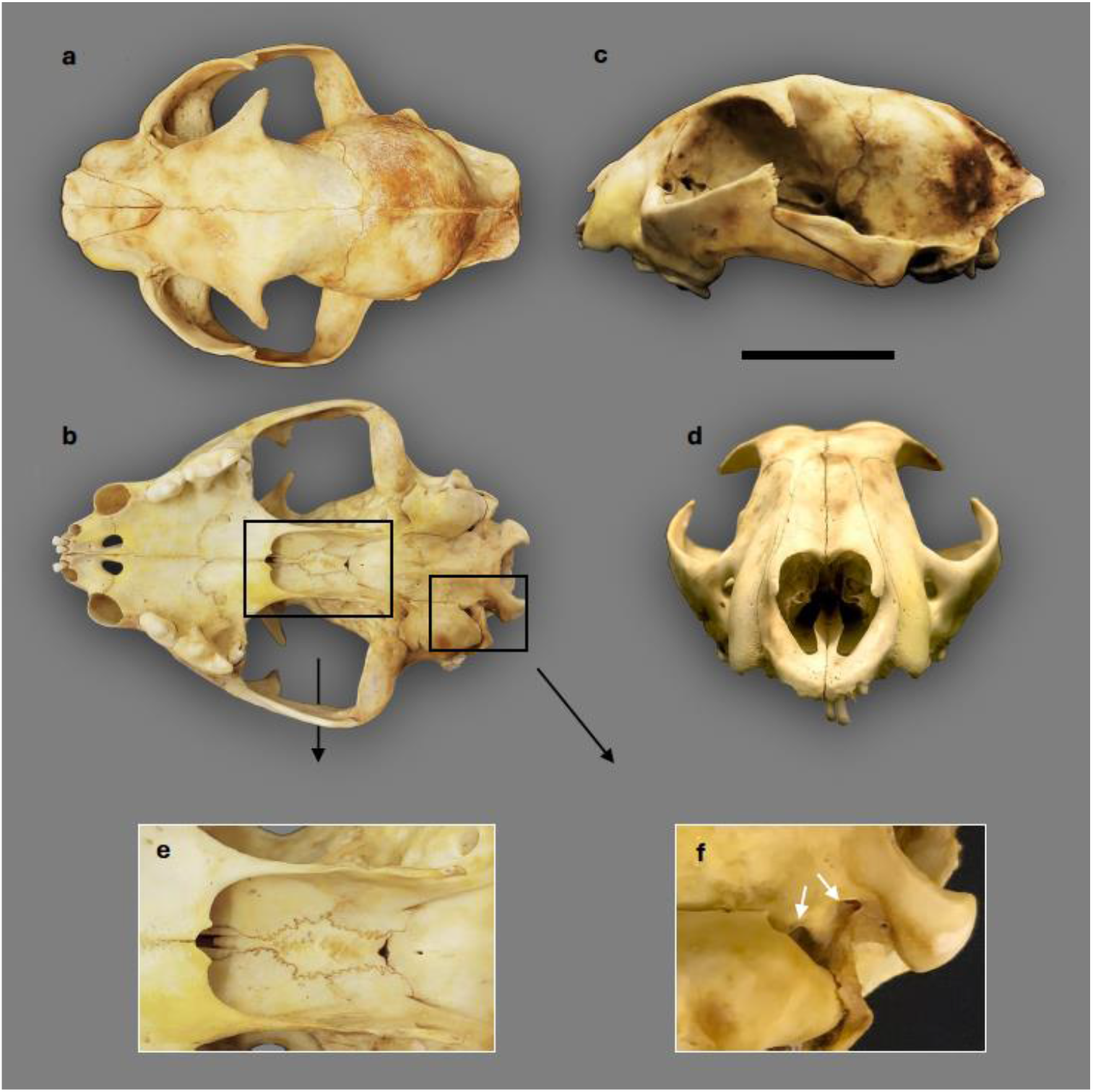
Skull of the Sima Topinoria specimen in a, dorsal; b, lateral; c, ventral; and d, frontal views; e, detail of the presphenoid synchondrosis; f, detail of the separation between hypoglossal and jugular foramina. Scale bar is 5 cm.

**Figure 6.**
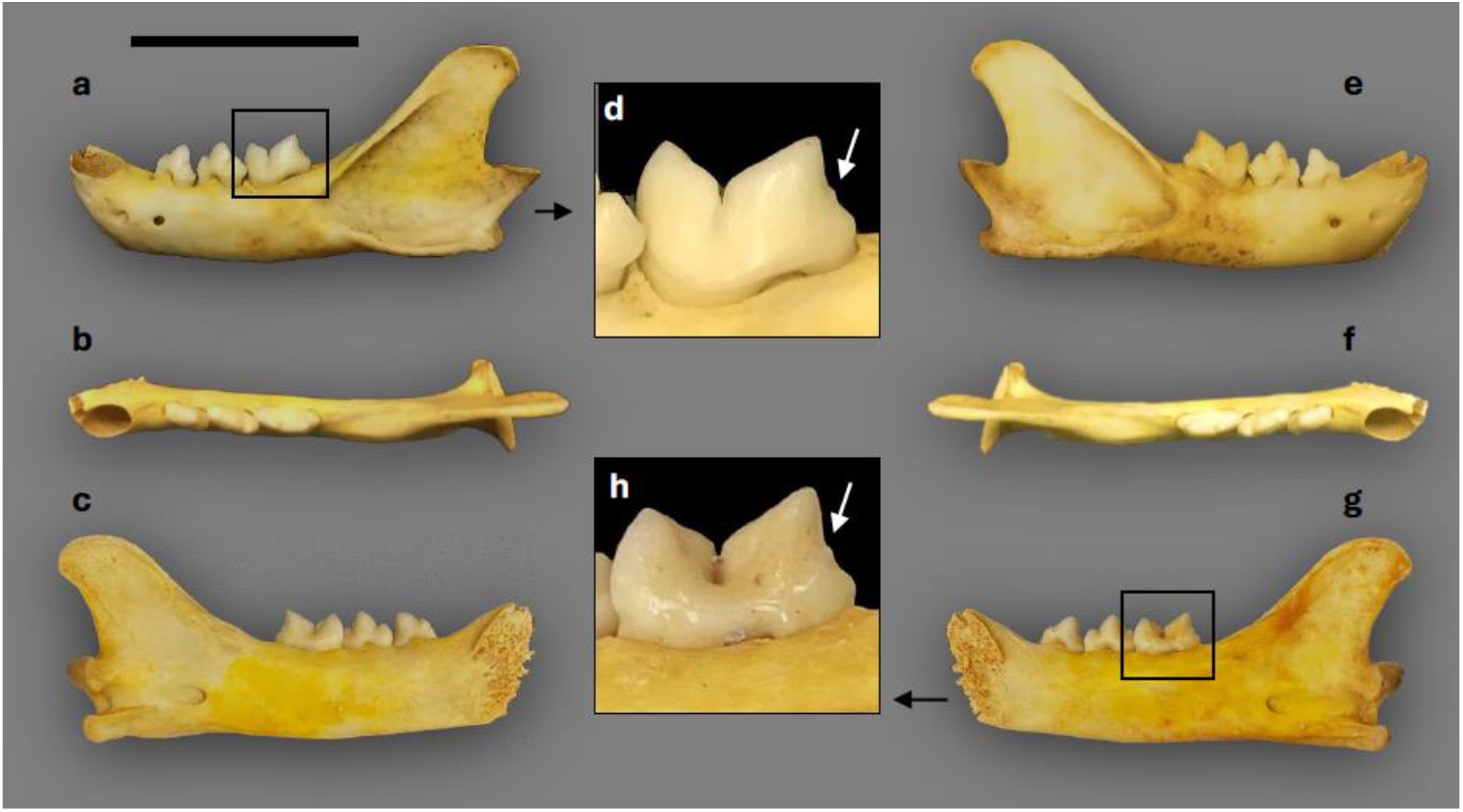
Left hemimandible of the Sima Topinoria specimen in 1a labial, 1b dorsal, and 1c lingual views. Detail in 1d shows the metaconid in the lower carnassial. Same for the right hemimandible 2a, 2b, 2c and 2d. Scale bar is 5 cm.

### 3.3. Metric data

Having a complete skeleton of an animal is essential for obtaining accurate metric data from each of its bones. This information provides reference measurements that can be used as a standard when analyzing fragmentary remains found in archaeopaleontological contexts. The anatomical integrity and excellent state of preservation of the specimen studied offer a unique opportunity to carry out a comprehensive metric study of the postcranial skeleton. The Supplementary Material 1 provides metric data for all the bone remains of the analyzed lynx.

### 3.4. Age, body mass, height at withers and sex

The skeleton of the Eurasian lynx from the Topinoria sinkhole exhibits a series of traits that identify it as an adult specimen, though not a senile one.

- All epiphyses of the long bones are fully fused, with no visible trace of the sutures.
- All vertebral discs are completely fused.
- All preserved teeth belong to permanent dentition and show no significant wear.
- The ossification of the presphenoid synchondrosis (a trait typical of the Eurasian lynx morphotype) is in an advanced stage (Fig. 5e). This corresponds to morphotype C (adult) as described by García-Perea (1996).
- There is no gap between the temporal crests at the level of the temporoparietal suture (Fig. 5a), also consistent with morphotype C (adult) in García-Perea (1996).
- The sagittal crest, although weakly developed, spans the entire temporal suture (Fig. 5a, 5b). This corresponds to morphotype D (adult) in García-Perea (1996).

Body mass can be estimated from skeletal measurements using allometric equations that relate specific bone dimensions to overall body size. One widely cited formula was proposed by Van Valkenburgh (1990) for felids in general, based on the condylobasal length of the skull: log_10_(body mass)=log_10_(basilar length) * 3.11 − 5.38. This model assumes a consistent allometric relationship between cranial size and body weight across different felid species. It is particularly valuable when only cranial elements are preserved.

According to this formula, the lynx from Torca de la Topinoria, whose condylobasal length is 140.1 mm, would weigh 19.7 kg. This value is similar to the average weight of modern males from Russia (19.6 kg, between 16.3 and 23.5), while the average weight of females is 17.3 kg (Sunquist and Sunquist, 2002), and Late Pleistocene specimens from Southern Europe, such as Grotte de l’Observatoire or Rèseau du Cèbèri (Mecozzi et al., 2021). The Sueve specimen (Nores, 1999), with a condylobasal length of 138 mm, could weigh 18.8 kg, while the Sima de Pagolusieta specimen in Navarre (Altuna, 1980), with a condylobasal length of 145 mm, would weigh 22 kg. A table with estimated body mass comparisons with published data from modern and ancient populations can be found in Supplementary Material 2.

In addition to body mass, shoulder height can also be estimated from postcranial elements. For lynx specifically, Gal et al. (2024) developed a regression model based on modern lynx skeletons that relate skull total length and long bones (humerus, radius, femur and tibia) lengths to estimated height at the withers. The equations use the logarithm of bone length to account for the nonlinear relationship between skull or limb proportions and body size. This allows for reasonably accurate height estimates even from fragmentary remains. Table 3 shows the calculation of the height at the withers of the Sima Topinoria lynx according to the equations proposed by Gal et al. (2024) for the skull and long bones, as well as these same calculations for the Pagolusieta and Sueve individuals. The height at the withers calculated from the total length of the skull is greater than that obtained from the long bones, which, for their part, show more homogeneous results.

**Table 3.** Estimated height at withers (HW) for Iberian *Lynx lynx* calculated from the total length (TL) of the skull and the limb long bones (Gal et al., 2024).

|  |  |  | Sima Topinoria |  | Pagolusieta | Sueve |
| --- | --- | --- | --- | --- | --- | --- |
| bone | Gal et al. (2024) equations | side | TL (mm) | HW (mm) | HW (mm) | HW (mm) |
| Skull | $\log HW = 1.309(\log TL) - 0.090$ | - | 156,6 | 606,7 | 644,5 | 578,5 |
| Humerus | $\log HW = 0.994(\log TL) + 0.499$ | left | 185 | 565,7 | 609,7 | - |
|  |  | right | 185 | 565,7 | - | - |
| Radius | $\log HW = 1.352(\log TL) - 0.290$ | left | 177,5 | 563,5 | - | - |
|  |  | right | 176 | 557,1 | - | - |
| Femur | $\log HW = 0.867(\log TL) + 0.725$ | left | 216 | 561,0 | 608,0 | - |
|  |  | right | 216 | 561,0 | - | - |
| Tibia | $\log HW = 1.234(\log TL) - 0.150$ | left | 222 | 556,4 | 592,2 | - |
|  |  | right | 221,5 | 554,9 | - | - |
| average long bones |  |  |  | 560,7 | 603,3 |  |
| st. dev. |  |  |  | 4,2 | 9,7 |  |

Based on cranial and postcranial estimations, the body mass of the Topinoria lynx is estimated as 19.7 kg and its height at withers around 56 cm, both within the range of modern Eurasian lynx males (see Supplementary Material 2). In this context, the Sueve specimen could have been a large female, akin to those from contemporary populations from higher latitudes and altitudes (Siberia, Tibet). Although it cannot be discarded the possibility of Sima Topinoria being a large female, as determining the sex of the individual is not straightforward. On the one hand, the baculum or penile bone that would clearly identify it as male has not been found. However, the penile bone in lynxes is practically vestigial, barely 5 mm long (Tumlinson, 1987; Larivière & Walton, 1997; Lavoie et al., 2019), so it may have gone unnoticed among the gravel and sediments on which the skeleton was found.

Some authors propose that size alone is the only determining factor in identifying the sex, since sexual dimorphism in non-social felines manifests itself only in body size and not in proportions or other characteristic features (Beltrán and Delibes, 1993; Bizhanova and Grachev, 2022). However, other authors propose that there is a certain difference between males and females in cranial proportions. In a metric study of lynxes and other felines, Sicurro and Oliveira (2011) found that in the Eurasian lynx, the postorbital constriction is the least dimorphic measurement, resulting in a greater relative width of the skull in females. This trait is probably due to the need for the neurocranium to maintain a functional size despite the smaller size of females, as observed in Pleistocene cave bears *Ursus spelaeus* (Grandal-d’Anglade & López González, 2005; Grandal d’Anglade, 2010). In the case of ST, we have been unable to analyse these measures due to a lack of comparative data.

### 3.5. Craniometric analysis

Due to limited comparable measurements, we selected the total skull length (TL) and bizygomatic breadth (ZB) for a comparative craniometric analysis. A bivariate plot (Fig. 7) was generated using the mean TL and ZB values to visualize cranial size and overall shape between populations and sexes.

**Figure 7.**
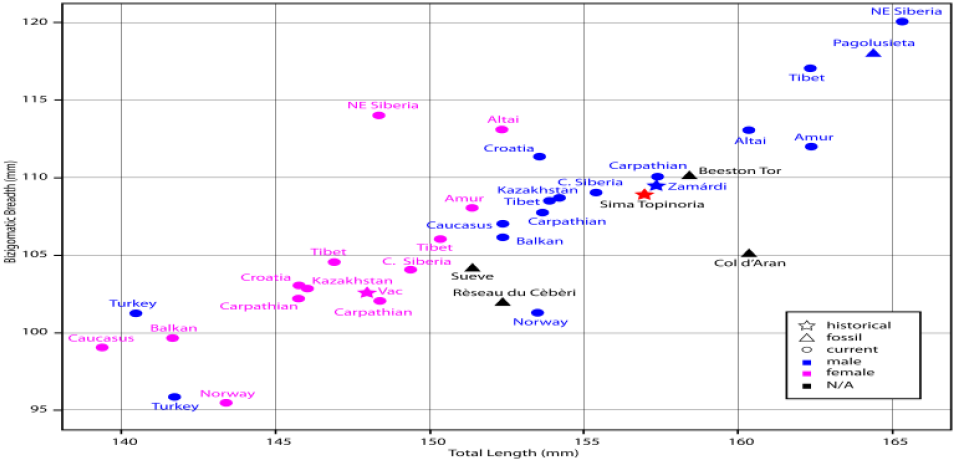
Bivariate diagram of total length and zygomatic breadth of the skull for the Sima Topinoria lynx and other Eurasian lynx populations (see Fig. 1) separated by chronological periods and sex.

Larger modern individuals, particularly those from colder ecosystems such as Siberian and Tibetan populations, score highest, whereas specimens from lower latitudes (e.g., Anatolia, the Caucasus, and the Balkans) exhibit smaller values, following a latitudinal size gradient consistent with Bergmann’s rule. This pattern is especially pronounced in the Anatolian lynx and aligns with previous studies describing smaller body size and a more specialized foraging ecology focused on brown hares rather than ungulates (Mengüllüoğlu et al., 2018; Soyumert et al., 2019). Northeastern Siberian, Altai, and Tibetan populations show the biggest skulls and broadest zygomatic arches among modern lynxes, though the latter display considerable intrapopulational variability. In contrast, Norwegian lynxes exhibit narrower skulls, a feature potentially reflecting postglacial bottlenecks, founder effects, or relaxed selection pressures. Despite their relatively gracile cranial morphology, their trophic ecology remains broadly similar to that of continental populations, with roe deer constituting the dietary core (Odden et al., 2006).

The Sima Topinoria specimen plots within the morphospace of Eurasian lynx males, closely overlapping with modern Carpathian and central Siberian samples, as well as with a male from medieval Hungary and the Late Pleistocene specimen from Beeston Tor (UK). Notably, the Sima Topinoria specimen plots far from other Iberian Eurasian lynx specimens, suggesting a reduction in size relative to the ∼2,570 y BP male from Pagolusieta (Altuna, 1980).**3.6. Pathologies**

The skeleton exhibits several pathological features. The proximal epiphysis of the right fibula is fused to the tibia. The second phalanx of the fourth digit on the right forelimb shows a pathological thickening, possibly resulting from an old fracture. The most notable pathological feature is that observed in the series of left ribs, affecting ribs two through eight. Both morphoscopic and radiographic analyses reveal posterior arch fractures in six of these ribs (Fig. 8a). Except for the first and second ribs, the rest (ribs 3 to 8) show fractures in the early stages of healing. In some cases, where there was no displacement of the fragments, a developing fracture callus with periosteal reaction is visible (Fig. 8b), characterized by poorly structured bone growth and marked radiolucency (Wedel & Galloway, 2014). In other ribs with fragment displacement, the periosteal reaction is more pronounced, and the callus has failed to unite the fragments (Ortner, 2003; Serrulla, 2015).

**Figure 8.**
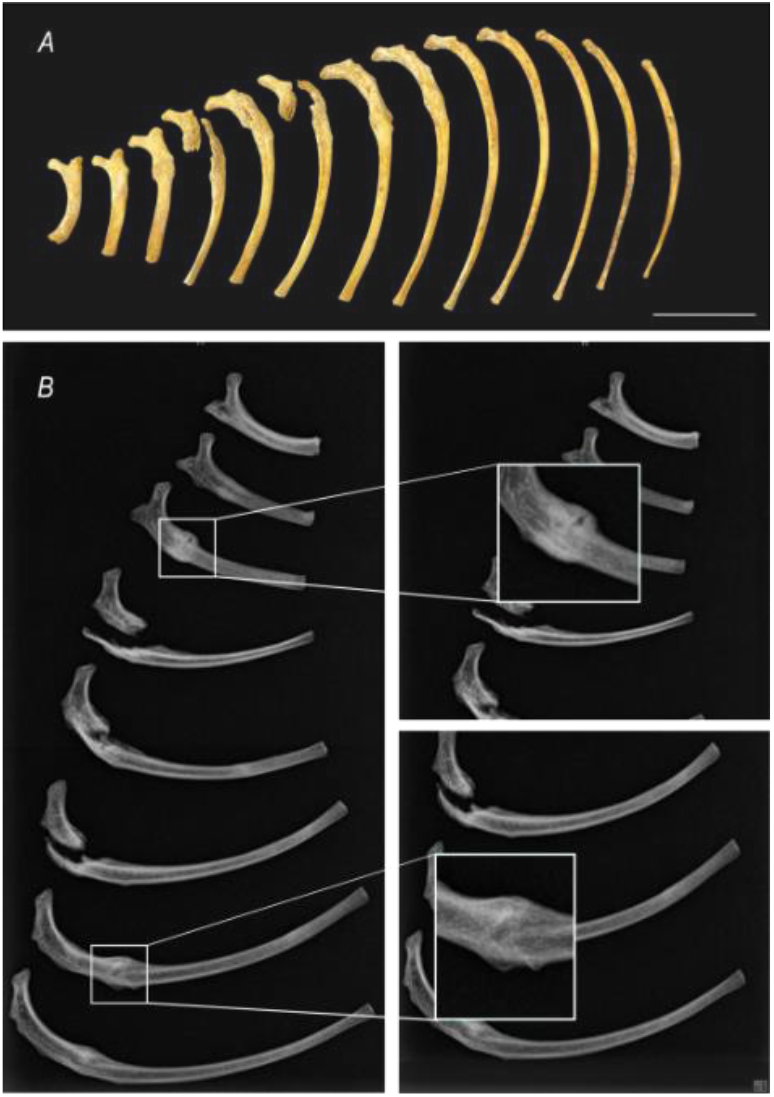
Thoracic pathologies A) Left rib cage showing pathologies consistent with fractures in multiple ribs. B) Radiograph of the left ribs affected by trauma. B1: Detail of the callus on the 3rd rib. B2: Detail of the callus on the 7th rib. Institute of Legal Medicine of Galicia. Scale bar: 5cm

Considering the distribution of the fractures and the healing stage of the fracture foci, we infer that the trauma resulted from a high-energy impact, and that the animal survived for approximately 10–20 days following the injury (Serrulla, 2015; Reichs, 1998). These fractures were likely caused by a fall into the cave.

### 3.7. Historical decline

A dramatic decline in the number of lynx records was recorded during the first half of the 19^th^ century throughout northern Spain (Table 4). In less than half a century, these records were reduced to one-tenth of their previous number, apparently indicating the disappearance of the lynx from the entire Pyrenean range. Municipalities with lynx presence show a widespread decline throughout the period in Asturias (PEF2x3 = 0.000003) and in the Basque Country (PEF2x2 = 0.003) between the first two periods. By the mid-20th century, geographical dictionaries recorded its presence in only a single locality in the Basque Country: Salvatierra (Álava).

**Table 4.**
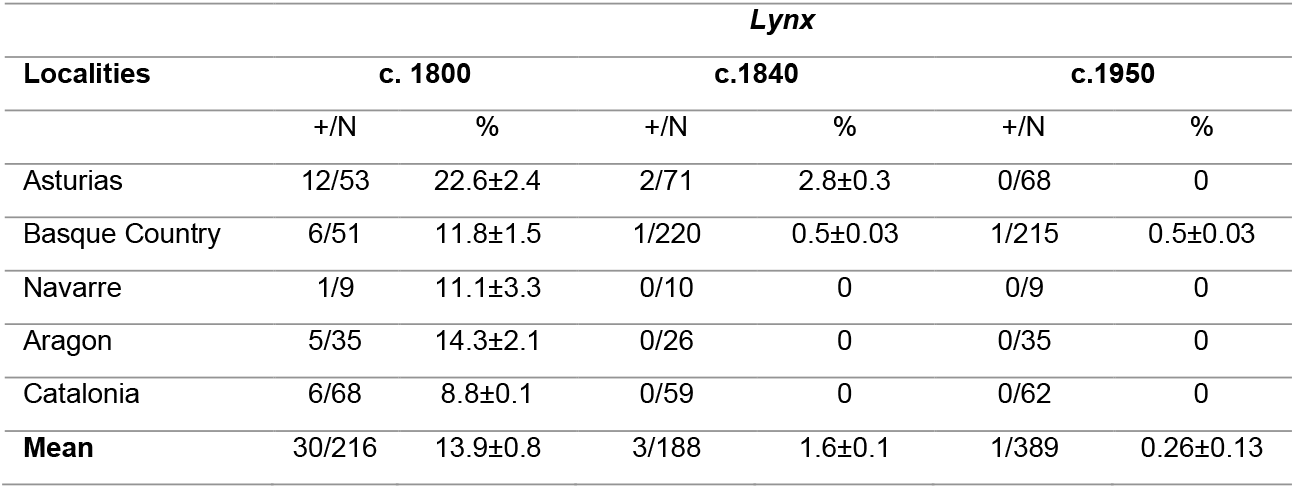
Reduction in lynx records in northern Spain from c. 1800 to c. 1950. Municipalities with presence (+) and total municipalities considered (N). Percentage ± SE (%).

| Localities | Lynx |  |  |  |  |  |
| --- | --- | --- | --- | --- | --- | --- |
|  | c. 1800 |  | c.1840 |  | c.1950 |  |
|  | +/N | % | +/N | % | +/N | % |
| Asturias | 12/53 | 22.6±2.4 | 2/71 | 2.8±0.3 | 0/68 | 0 |
| Basque Country | 6/51 | 11.8±1.5 | 1/220 | 0.5±0.03 | 1/215 | 0.5±0.03 |
| Navarre | 1/9 | 11.1±3.3 | 0/10 | 0 | 0/9 | 0 |
| Aragon | 5/35 | 14.3±2.1 | 0/26 | 0 | 0/35 | 0 |
| Catalonia | 6/68 | 8.8±0.1 | 0/59 | 0 | 0/62 | 0 |
| **Mean** | 30/216 | 13.9±0.8 | 3/188 | 1.6±0.1 | 1/389 | 0.26±0.13 |

## 4. DISCUSSION

### 4.1. Historical context

The confirmation of the lynx from Torca de la Topinoria provides physical evidence that the Eurasian lynx (*Lynx lynx*) persisted in the Iberian Peninsula until relatively recent times, and it reinforces the coherence of the hypothesis that the vernacular name “lobo cerval” referred to *Lynx lynx*, a larger species found in Atlantic environments, whereas “gato cerval” referred to the smaller, Mediterranean Lynx pardinus (Clavero & Delibes, 2013).

Records of the lobo cerval are abundant throughout the 18th century and early 19th century, from northern Portugal (Carvalho, 1706) and various points in the Cantabrian Mountains (Sarmiento, 1760; Bowles, 1775) to the Pyrenees (Assó, 1784). Numerous studies have also examined its persecution in Asturias (Torrente, 1999) and the Basque Country (López de Guereñu, 1957; Fernández & Ruiz de Azúa, 2003; Garayo, 2021).

The decline of the lynx during the first half of the 19th century is consistent with the effects of institutional campaigns of poisoning with nux vomica and strychnine aimed at eradicating vermin, initiated in 1829. These campaigns have likewise been linked to a significant reduction, fragmentation, and local extinction of the wolf in Spain (Nores & López-Bao, 2022), a carnivore more abundant and ecologically plastic than the lynx. During the same period, similar effects were observed in other scavenging species, such as wild boar in Asturias (Nores et al., 1995).

It is often assumed that generalist carnivores, reputed scavengers such as wolves and foxes, are more sensitive to poisoned baits than specialized carnivores such as the lynx. However, it has been shown that lynxes use carrion as an important dietary supplement (Odden et al., 2006; Premier et al., 2021), making them facultative scavengers (King et al., 2015), which also explains their sensitivity to poison placed in carcasses.

The absence of lynx records from the Pyrenees in geographical dictionaries from the mid-19th century reflects a widespread decline, as also noted by Trutat (1878) for the French side, but not a total extinction, since Martínez Reguera (1881) observed the species in the province of Huesca (Aragón), along with other less explicit records in Navarre (Ibarlucea, 1886).

### 4.2. Environmental Context: The Picos de Europa as a Potential Refugium for the Eurasian Lynx

The recent dating of the Eurasian lynx remains from Sima Topinoria suggests that relict populations of this species may have persisted in the Iberian Peninsula until much more recent times than previously assumed. In this context, the Picos de Europa National Park provides an especially suitable environmental setting to explore this hypothesis.

Established in 1918 as Spain’s first national park, the Picos de Europa are characterized by their striking limestone landscapes and remarkable biological diversity. Initially limited to the Western Massif, covering around 17,000 hectares, the park was progressively expanded to encompass all three massifs that give the park its name, currently extending over approximately 67,000 hectares. In 2003, it was designated a UNESCO Biosphere Reserve.

Its proximity to the Cantabrian coast (about 30 km) and the marked altitudinal range, with peaks exceeding 2,500 m a.s.l., create substantial ecological heterogeneity. The combination of deep valleys and gorges with these altitudinal gradients allows for the coexistence of Atlantic, Mediterranean and boreo-alpine taxa, as well as a high level of endemism. If relict lynx populations were to have survived in any part of the Peninsula in recent times, this park offers especially favourable conditions.

The landscape forms a mosaic of ecosystems, among which mature forests stand out, covering approximately 25% of the park’s surface (Fernández-Prieto & Bueno, 2013; González-García et al., 2024). These include beech forests, oak woodlands, Eurosiberian mixed forests and subalpine formations, all potentially suitable habitats for the Eurasian lynx, which requires dense forest cover, as well as rugged terrain, higher slopes or low-altitude mountain pastures (Conc et al., 2024; Nagl et al., 2022).

### 4.3. Taxonomic and chronological contextualization

The diagnostic cranial and dental traits of a novel *Lynx* skeleton discovered in the Iberian Peninsula Topinoria specimen unequivocally identify them as *Lynx lynx*. The presence of V-shaped temporal crests separated hypoglossal and jugular foramina, a distinct metaconid on the lower carnassial and its high body mass and overall size collectively exclude *L. pardinus* (García-Perea, 1985, 1996; Kitchener et al., 2017; Mecozzi et al., 2021).

The calibrated age of the Topinoria lynx (1729–1808 cal AD; 51.6% probability) identifies it as the latest confirmed record of *Lynx lynx* in the Iberian Peninsula, and at least two centuries after the youngest previously known Eurasian lynx remains from mid-15^th^ to mid-16^th^ centuries (Rodríguez-Varela et al., 2016) evidencing the persistence of the species in northern Spain into the 18^th^-19^th^ century, much later than formerly assumed.

This finding aligns with scattered historical reports of large felids in Cantabrian and Pyrenean regions into early modern times (Altuna, 1980; Delibes, 1983; Clavero & Delibes, 2013), though such accounts have often been difficult to attribute reliably to *Lynx lynx* rather than *L. pardinus*. The Topinoria specimen provides the first direct material and chronological evidence confirming that the Eurasian lynx survived in the Cantabrian region into modern times. Its survival likely reflects the persistence of suitable forested habitats in high-altitude refugia, as lowland environments became increasingly fragmented by agricultural and industrial expansion (Carrión et al., 2010).

The estimated body mass of 19.7 kg and shoulder height of ca. 56 cm place the Topinoria lynx within the range of modern adult males from Central and Eastern Europe (Sunquist & Sunquist, 2002; Červený & Koubek, 2000). Accordingly, the craniometric analysis positions the Topinoria specimen within the morphometric range of Western and Central Eurasian *Lynx lynx* males, clustering particularly with modern Carpathian specimens from Czechia (Červený & Koubek, 2000), a medieval male from Hungary (Gal et al., 2022, 2024), and contemporary males from Central Russia (Stroganov, 1969).

Beyond this general affinity, our results reveal significant cranial shape variation among Iberian *L. lynx* specimens from different temporal contexts. The protohistoric specimen from Pagolusieta (ca. 2570 yBP) exhibits a robust and elongated skull, with broad zygomatic arches and a pronounced sagittal crest, indicating a generally larger and more massive morphology. Similar robust cranial morphologies have been documented in fossil Eurasian lynxes from the Pyrenees (Castaños, 1987) and are consistent with patterns observed in other Quaternary mammalian assemblages from the Cantabrian region (Fidalgo et al., 2025, but see also García-Vázquez, 2025).

These differences suggest a temporal trend of body size reduction in *L. lynx* from the Pleistocene to the present, a pattern also described for *L. pardinus* (Kurtén, 1968; Mecozzi et al., 2021). Given that the Pagolusieta specimen dates to the first millennium BCE, this trend would imply a significant reduction in body size since protohistoric times, rather than being restricted to strictly postglacial processes. Comparable size decreases have been reported in other large carnivores from the Iberian Quaternary sequence (Castaños, 1987) and the Cantabrian Mountains (Fidalgo et al., 2025; García Vázquez, 2025).

This pattern is further supported by the scarce postcranial evidence available from Spain, including the larger metatarsal II from Rascaño (11917 yBP) and Pagolusieta (2750 yBP) (Altuna, 1980) (Supplementary Material 2), as well as metatarsal IV elements from Pagolusieta (Altuna, 1980), Santimamiñe and the French sites of Moustayous, Labastide and Asson (Castaños, 1987). Together, these data point to a gradual reduction in skeletal size through time.

In this context, the smaller and more gracile skull of the Sima Topinoria specimen, as well as its slightly reduced cranial metrics compared to protohistoric Iberian males such as Pagolusieta, may represent a later, geographically isolated lineage. Such morphological shifts are consistent with long-term confinement and environmental constraints in the Cantabrian Mountains, potentially associated with reduced genetic diversity or limited prey availability. Similar processes of localized adaptation and size reduction have been proposed for isolated or relict populations of large carnivores (López-Bao et al., 2010).

This interpretation is further reinforced by the chronological context of the Topinoria specimen, corresponding to a period of intensified human disturbance, industrial expansion and habitat degradation (Carrión et al., 2010), direct persecution (Villalpando-Moreno, 2020), and prey depletion linked to the widespread use of firearms (García-Vázquez, 2025). If we compare the biometric data of the Sima Topinoria lynx with the historical records transcribed by Nores et al (2015), it is lighter than the one hunted in Motriku (Basque Country) in 1762, which weighed ‘fifty pounds (23 kg) or more’. Its size is similar to that of the specimen killed in San Pedro de los Montes (Ponferrada, León) mentioned by Fray Martín Sarmiento in 1760 and somewhat smaller than those mentioned by Martínez Marina in Langreo (Asturias) in 1802, which are believed to have been ‘about a yard (84 cm) tall’ (Nores et al., 2015). All these factors likely exacerbated ecological stress in an already fragmented population, contributing to the final decline and eventual extinction of the species in Spain.

Although other striking measurements were encountered in this specimen, such as the abnormally broad interorbital space (see Supplementary Material 1), these cannot be interpreted with the current data due to the lack of published material. Overall, our findings highlight the urgent need for a comprehensive metric reassessment of ancient Eurasian lynxes. The scarcity of available data and inconsistencies in measurement protocols currently limits the morphometric study of this species.

### 4.4. Pathological Evidence and Circumstances of Death

The skeletal pathologies observed in the Topinoria individual, particularly multiple rib fractures in early healing stages, suggest that the animal sustained a severe traumatic impact from which it survived for at least one to two weeks. The absence of predation or hunting marks, together with the location of the find within a vertical shaft, indicates that the lynx likely fell accidentally into the sinkhole and was subsequently unable to escape. Similar taphonomic scenarios have been described for other Pleistocene and Holocene felids in karstic contexts of northern Spain (Altuna, 1980; Villaluenga, 2016) and Southern France (Fosse et al., 2021). Additionally, other skeletal pathologies provide further insights into the individual’s life history. The left scapula presents a fracture consistent with the forces generated during the fall into the shaft, while the deformity observed in the second phalanx of the fourth manus digit (specimen no. 129) correspond to an antemortem, fully healed injury, demonstrating that the animal survived an earlier trauma long before the fall. Comparable cases of long-term survival following serious damage have been reported in a Pleistocene lynx from Taurida Cave, Crimea, which endured and recovered from major trauma (Serdyuk et al., 2024), as well as in modern Iberian lynxes that survived severe injuries without compromising reproductive capacity (García-Perea, 2000). Together, these observations highlight the resilience of lynxes and support the interpretation that the Topinoria lynx recovered from significant injuries prior to its accidental death.

## 5. CONCLUSIONS

The lynx skeleton recovered from the Topinoria sinkhole has been identified as a Eurasian lynx (*Lynx lynx* Linnaeus, 1758), based on several cranial features that distinguish it from the Iberian lynx. It is an almost complete skeleton, found in anatomical position and at considerable depth within the sinkhole, raising questions about how the animal accessed the site. At some undetermined point prior to the discovery of the remains by cavers from the Flash and Tracalet speleology groups, the skull and mandibles were moved, and the four canines were likely removed, as they were not recovered despite the highly meticulous excavation of the site.

Radiocarbon dating (^14^C) indicates that the animal most likely lived during the late 18^th^ to early 19^th^ centuries. The dates obtained are more recent than those of the latest known Eurasian lynx specimen in the Iberian Peninsula, dated to approximately 412 years BP. Thus, the Topinoria lynx represents the most recent known remains of *Lynx lynx* from the Iberian Peninsula.

The individual was an adult, though not senile, and its body mass is estimated at around 20 kg. The skull is similar in size to that of adult males from present-day populations in Central Europe and Central Asia, which suggests it may have been a male. However, the absence of other contemporary specimens from the Cantabrian Mountains or the Pyrenees limits the possibility of a meaningful comparative metric analysis, except in relation to current extra-peninsular populations. Among those, the Topinoria lynx displays a relatively large skull size for male individuals, albeit smaller when compared to a medieval male from Zamárdi (Hungary) and older Holocene specimens from the Cantabrian region (Pagolusieta, Biscay), suggesting a recent size reduction, as observed for other mammals of the Cantabrian Mountains (Fidalgo et al., 2025).

This finding not only lends strong support to the hypothesis that the Eurasian lynx survived in this region into very recent times, but it also brings new relevance to long-standing oral traditions in northern Spain. In the Cantabrian region, local folklore includes recurring accounts of a mysterious animal known as the *lobo cerval*, described as a large, elusive feline with a spotted or mottled coat. These accounts, preserved through generations, may indeed reflect cultural memory of the Eurasian lynx, a species whose presence had not been archaeologically documented in the area in historical times, until now.

From a taxonomic and methodological perspective, the study of this skeleton is also important due to the potential for misidentification of lynx remains in Holocene contexts. Given the likely spatiotemporal overlap of *Lynx lynx* and *Lynx pardinus* in northern Iberia during certain climatic phases, the clear morphological and biometric distinction between the two becomes essential. In fragmentary or poorly preserved remains, species-level identification can be challenging, and the existence of a robust osteometric reference for historical *Lynx lynx* in Iberia can significantly enhance the accuracy of future identifications.

The discovery of the Topinoria lynx substantially extends the known temporal range of *L. lynx* within the Iberian Peninsula and refines our understanding of its final extinction dynamics. The specimen demonstrates that isolated populations persisted in the Cantabrian Mountains well into the 18th–19th centuries and confirms that both the Iberian lynx (*Lynx pardinus*) and the Eurasian lynx coexisted on the peninsula until that period. This finding highlights the species’ status as autochthonous to the Iberian Peninsula and indicates that suitable ecological conditions for both lynxes persisted in northern Spain well into the modern era (Gil-Sánchez & McCain, 2011). The Topinoria lynx provides critical historical context for future studies of lynx ecology, biogeography, and species interactions, as well as for future research on the ecological and conservation dynamics of the region, including possible reintroduction proposals.

Ultimately, the study of this skeleton not only fills a critical gap in the recent faunal history of the Iberian Peninsula but also provides essential reference data for improving the resolution of future taxonomic and ecological research on lynx populations in Europe.

## Acknowledgements

This work was promoted and funded by the Picos de Europa National Park, with the authorisation of the General Directorate of Cultural Heritage and Historical Memory of the Government of Cantabria. The authors would like to thank the members of the Flash Caving Group who collected the lynx remains in the Sima Topinoria along J.A. Martín Otero: Jorge Mateos de la Fuente, José Luis Izquierdo Moreno, Juan Bueno Gabaldón, Eduardo Tomás Mezquida, as well as Belén Hernández Fernández on the external team. We would also like to thank the Park Rangers Miguel A. Rabanal and Jorge García for their collaboration in the field campaign, and also the shepherd Braulio Roiz and his family for their hospitality and knowledge of the area.

