## Supplementary Material 1 for "The Last Iberian Record of Eurasian Lynx (*Lynx lynx*): Osteometry and Historical Implications of the Lynx from Sima Topinoria (Cantabria, Spain)"

#### Appendix I

Measurements of the skeletal remains of the lynx from Sima Topinoria

NOTES:
The plates show the paired bones of the right side.
Values in italics denote measurements that are not comparable because they are pathological.
An asterisk marks questionable measurements due to worn remains.

| CRANIUM (Plate 1) | | | | | |
|---|---|---|---|---|---|
| code | description | Sima Topinoria | | Sueve | |
| 1 | Total length: Acrocranion - Prosthion | 156.6 | | 151.0 | |
| 2 | Condylobasal length: aboral border of the occipital condyles - Prosthion | 140.1 | | 138.0 | |
| 3 | Basal length: Basion - Prosthion | 129.5 | | | |
| 4 | Basicranial axis: Basion - Synsphenion | 58.0 | | | |
| 5 | Basifacial axis: Synsphenion - Prosthion | 82.4 | | 85.0 | |
| 6 | Median palatal length: Staphylion - Prosthion | 58.7 | | | |
| 7 | Zygomatic breadth: Zygion - Zygion | 108.9 | | 104.0 | |
| 8 | Greatest mastoid breadth: Otion - Otion | 66.7 | | 66.2 | |
| 9 | Frontal breadth: Ectorbitale - Ectorbitale | 69.9 | | | |
| 10 | Least breadth between the orbits: Entorbitale - Ectorbitale | 45.6 | | 39.0 | |
| 11 | Greatest neurocranium breadth: Euryon - Euryon | 59.2 | | | |
| 12 | Height of the occipital triangle: Acrocranion - Basion | 47.7 | | | |
| 13 | Upper neurocranium length: Acrocranion - Frontal midpoint | 87.4 | | 90.0 | |
| 14 | Facial length: Frontal midpoint - Prosthion | 79.3 | | 81.0 | |
| 15 | Neurocranium length: Basion - Nasion. | 113.7 | | 105.0 | |
| 16 | Viscerocranium length: Nasion - Prosthion | 49.0 | | 64.0 | |
| 17 | Greatest palatal breadth | 59.0 | | 65.0 | |
| 18 | Breadth at the canine alveoli | 42.6 | | 40.2 | |
| 19 | Least breadth of the postorbital constriction | 39.7 | | | |
| 20 | Facial breadth between the infraorbital foramina | 45.6 | | | |
| 21 | Lateral length of snout: oral border of the orbit - Prosthion | 47.9 | | 45.0 | |
| 22 | Length of the cheekteeh row | 31.0 | 30.3 | | |
| 23 | Length of the premolar row | 5.1 | 5.2 | 30.0 | 30.0 |
| 24 | Length of diastema | 29.8 | 29.0 | 6.0 | 6.0 |
| 25 | Greatest diameter of the auditory bulla | 23.5 | | 30.0 | |
| 26 | Least diameter of the auditory bulla | 15.4 | | 15.0 | |
| 27 | Greatest breadth of the foramen magnum | 19.8 | | | |
| 28 | Height of the foramen magnum: Basion - Opisthion | 17.7 | | | |
| 29 | Greatest inner height of the orbit | 39.2 | | 30.0 | |
| 30 | Greatest inner length of the orbit: Ectorbitale - Ectorbitale | 38.4 | | 45.0 | |
| 31 | Transversal breadth of canine alveolus | 8.1 | 8.7 | | |
| 32 | Anteroposterior breadth of canine alveolus | 10.9 | 11.4 | | |
| **Upper cheek teeth** | | **dex** | **sin** | **dex** | **sin** |
| 1 | P3 length | 12.5 | 11.7 | | |
| 2 | P3 breadth | 6.4 | 6.4 | | |
| 1 | P4 length | 16.2 | 16.7 | | |
| 2 | P4 breadth | 9.9 | 10.0 | | |
| 1 | M1 length | 9.5 | 9.4 | | |
| 2 | M1 breadth | 4.3 | 4.4 | | |

| MANDIBLE (Plate 1) | | | | | |
|---|---|---|---|---|---|
| | | Sima Topinoria | | Sueve | |
| code | description | dex | sin | dex | sin |
| 1 | Total length: condyloid process - Infradentale | 104.9 | 105.2 | 102.5 | |
| 2 | Length from the mandibular notch - Infradentale | 99.1 | 99.1 | 92.0 | |
| 3 | Height of the vertical ramus: angular process - coronoid process | 47.1 | 47.6 | 50.0 | |
| 4 | Height of the horizontal ramus behind M1 | 20.8 | 22.5 | 21.3 | |
| 5 | Height of the horizontal ramus in front of P3 | 19.3 | 18.6 | 20.4 | |
| 6 | Length of the cheekteeth row | 37.2 | 38.3 | 37.0 | |
| 7 | L diastema | 7.7 | 7.9 | 8.8 | |
| 8 | Breadth of condyloid process | 25.5 | 23.8 | | |
| 9 | Height of condyloid process | 7.9 | 7.7 | | |
| 10 | Transversal breadth of canine alveolus | 8.6 | 8.9 | | |
| 11 | Anteroposterior breadth of canine alveolus | 10.7 | 10.9 | 10.0 | |
| **Lower cheek teeth** | | **dex** | **sin** | **dex** | **sin** |
| 1 | p3 length | 9.9 | 10.2 | | |
| 2 | p3 breadth | 5.1 | 5.3 | | |
| 1 | p4 length | 11.9 | 12.6 | | |
| 2 | p4 breadth | 5.9 | 6.0 | | |
| 1 | m1 length | 15.2 | 15.3 | | |
| 2 | m1 breadth | 7.0 | 6.7 | | |

### Plate 1
### SKULL and JAW

**SKULL**

1 Total length: Acrocranion - Prosthion
2 Condylobasal length: aboral border of the occipital condyles - Prosthion
3 Basal length: Basion - Prosthion
4 Basicranial axis: Basion - Synsphenion
5 Basifacial axis: Synsphenion - Prosthion
6 Median palatal length: Staphylion - Prosthion
7 Zygomatic breadth: Zygion - Zygion
8 Greatest mastoid breadth: Otion - Otion
9 Frontal breadth: Ectorbitale - Ectorbitale
10 Least breadth between the orbits: Entorbitale - Ectorbitale
11 Greatest neurocranium breadth: Euryon - Euryon
12 Height of the occipital triangle: Acrocranion - Basion
13 Upper neurocranium length: Acrocranion - Frontal midpoint
14 Facial length: Frontal midpoint - Prosthion
15 Neurocranium length: Basion - Nasion.
16 Viscerocranium length: Nasion - Prosthion
17 Greatest palatal breadth
18 Breadth at the canine alveoli
19 Least breadth of the postorbital constriction
20 Facial breadth between the infraorbital foramina
21 Lateral length of snout: oral border of the orbit - Prosthion
22 Length of the cheekteeh row
23 Length of the premolar row
24 Length of diastema
25 Greatest diameter of the auditory bulla
26 Least diamenter of the auditory bulla
27 Greatest breadth of the foramen magnum
28 Height of the foramen magnum: Basion - Opisthion
29 Greatest inner height of the orbit
30 Greatest inner length of the orbit: Ectorbitale - Ectorbitale
31 Transversal breadth of canine alveolus
32 Anteroposterior breadth of canine alveolus

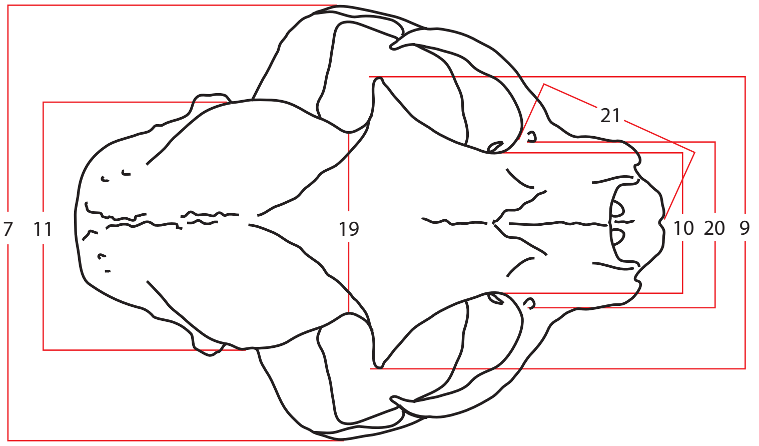

**JAW**

1 Total length: condyloid process - Infradentale
2 Length from the mandibular notch - Infradentale
3 Heigth of the vertical ramus: angular - coronoid processes
4 Height of the horizontal ramus behind M1
5 Height of the horizontal ramus in front of P3
6 Length of the cheekteeth row
7 Length of diastema
8 Breadth of condyloid process
9 Height of condyloid process
10 Transversal breadth of canine alveolus
11 Anteroposterior breadth of canine alveolus

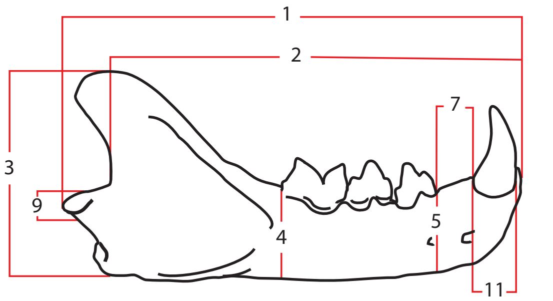

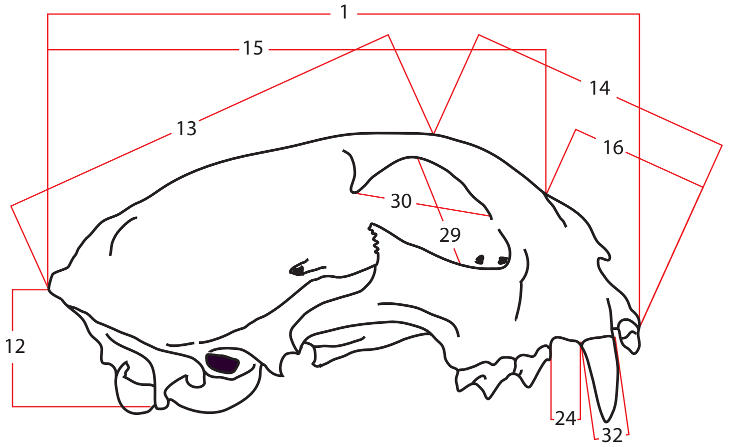

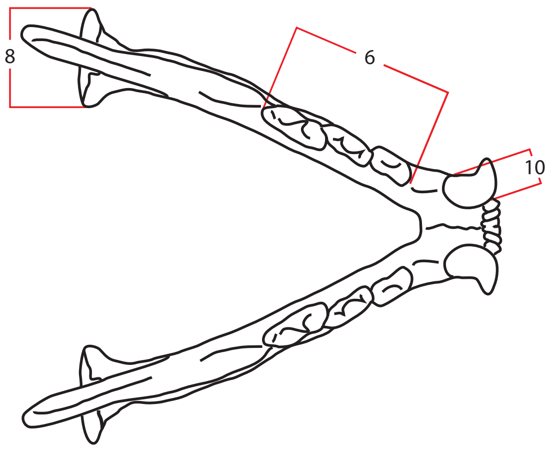

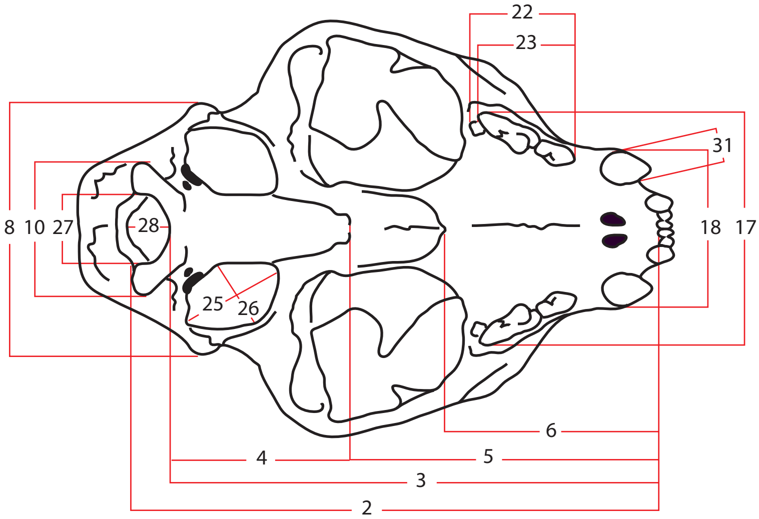

**CHEEK TEETH (P3, P4, M1, p3, p4, m1)**

1 Total length
2 Breadth

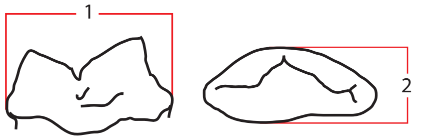

**AXIAL ELEMENTS and GIRDLES (Plate 2)**

| ATLAS | | |
|---|---|---|
| **code** | **description** | |
| 1 | Greatest breadth over the wings | 67.7 |
| 2 | Greatest length | 36.0 |
| 3 | Greatest breadth of the cranial articular surface | 38.2 |
| 4 | Greatest breadth of the caudal articular surface | 32.25 |
| 5 | Length of the dorsal arch | 15.2 |
| 6 | Height | 22.7 |
| **AXIS** | | |
| 1 | Greatest length in the region of the corpus including the dens | 43.6 |
| 2 | Greatest length of the arch including the caudal articular process | 40.1 |
| 3 | Greatest breadth of the cranial articular surface | 29.1 |
| 5 | Greatest breadth across the caudal articular process | 29.0 |
| 4 | Greatest breadth across the transversal process | 24.9 |
| 5 | Smallest breadth of the vertebra | 21.5 |
| 6 | Greatest breadth of the caudal articular surface | 18.8 |
| 7 | Greatest height | 34.2 |
| **SACRUM** | | |
| 1 | Greatest length on the ventral side | 56.2 |
| 2 | Physiological length | 46.8 |
| 3 | Greatest breadth across the wings | 43.6 |
| 4 | Greatest breadth of the cranial articular surface | 25.8 |
| 5 | Greatest height of the cranial articular surface | 13.1 |

| SCAPULA | | | |
|---|---|---|---|
| **code** | **description** | **dex** | **sin** |
| 1 | Height along the spine | 127.9 | 125.9 |
| 2 | Diagonal greatest height | 137.4 | 135.8 |
| 3 | Smallest length of the neck of the scapula | 25.3 | 25.2 |
| 4 | Greatest length of the glenoid process | 28.4 | 28.9 |
| 5 | Length of the glenoid cavity | 26.3 | 24.2 |
| 6 | Breadth of the glenoid cavity | 17.2 | 19.5 |

| PELVIS | | | |
|---|---|---|---|
| **code** | **description** | **dex** | **sin** |
| 1 | Greatest length of the half | 148.2 | 148.7 |
| 2 | Length of the acetabulum on the rim | 21.3 | 21.2 |
| 3 | Length of the symphysis (if fused) | 52.7 | |
| 4 | Smallest height of the shaft of ilium | 22.4 | 22.7 |
| 5 | Smallest breadth of the shaft of ilium | 9.6 | 9.4 |
| 6 | Inner length of the obturator foramen | 37.6 | 37.8 |
| 7 | Greatest breadth across the iliac crests | 66.1 | |
| 8 | Greatest breadth across the acetabula | 63.3 | |
| 9 | Greatest breadth across the ischiatic tuberosity | 75.3 | |
| 10 | Smallest breadth across the bodies of the ischia | 53.1 | |

### Plate 2
### AXIAL SKELETON and GIRDLES

**ATLAS**
1 Greatest breadth over the wings
2 Greatest length
3 Greatest breadth of the cranial articular surface
4 Greatest breadth of the caudal articular surface
5 Length of the dorsal arch
6 Height

**AXIS**
1 Greatest length in the region of the corpus including the dens
2 Greatest length of the arch including the caudal articular process
3 Greatest breadth of the cranial articular surface
5 Greatest breadth across the caudal articular process
4 Greatest breadth across the transversal process
5 Smallest breadth of the vertebra
6 Greatest breadth of the caudal articular surface
7 Greatest height

**SCAPULA**
1 Heigth along the spine
2 Diagonal greatest height
3 Smallest length of the neck of the scapula
4 Greatest length of the glenoid process
5 Length of the glenoid cavity
6 Breadth of the glenoid cavity

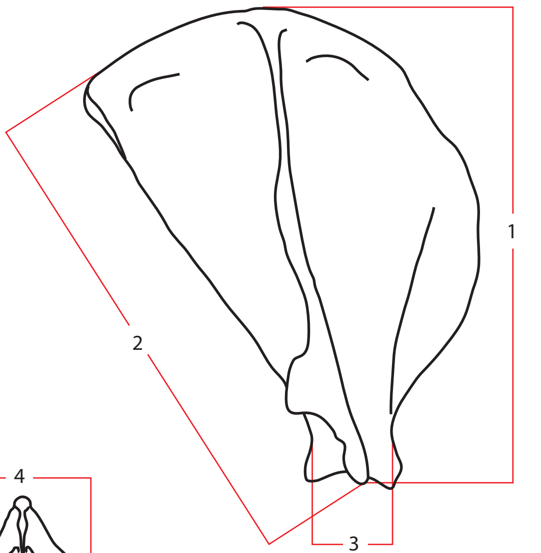

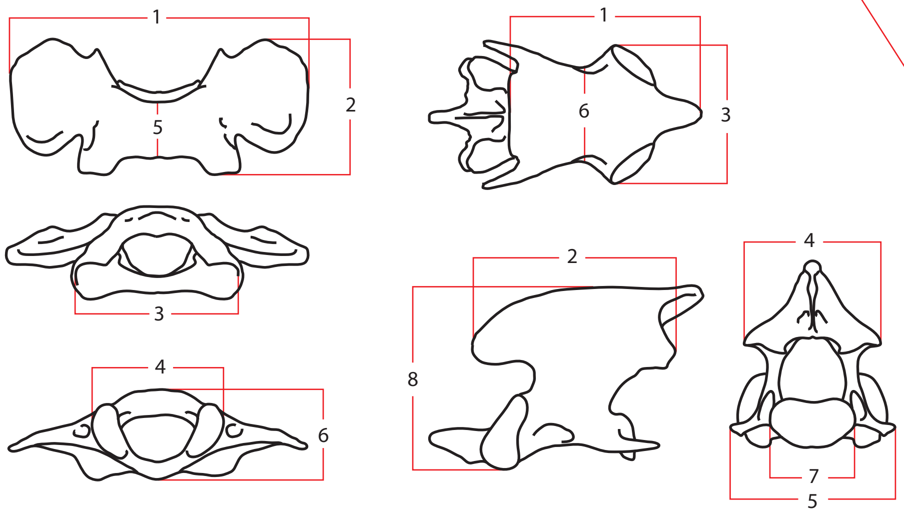

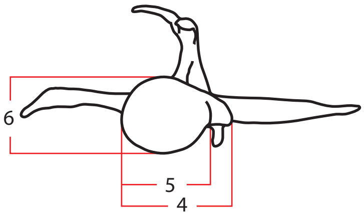

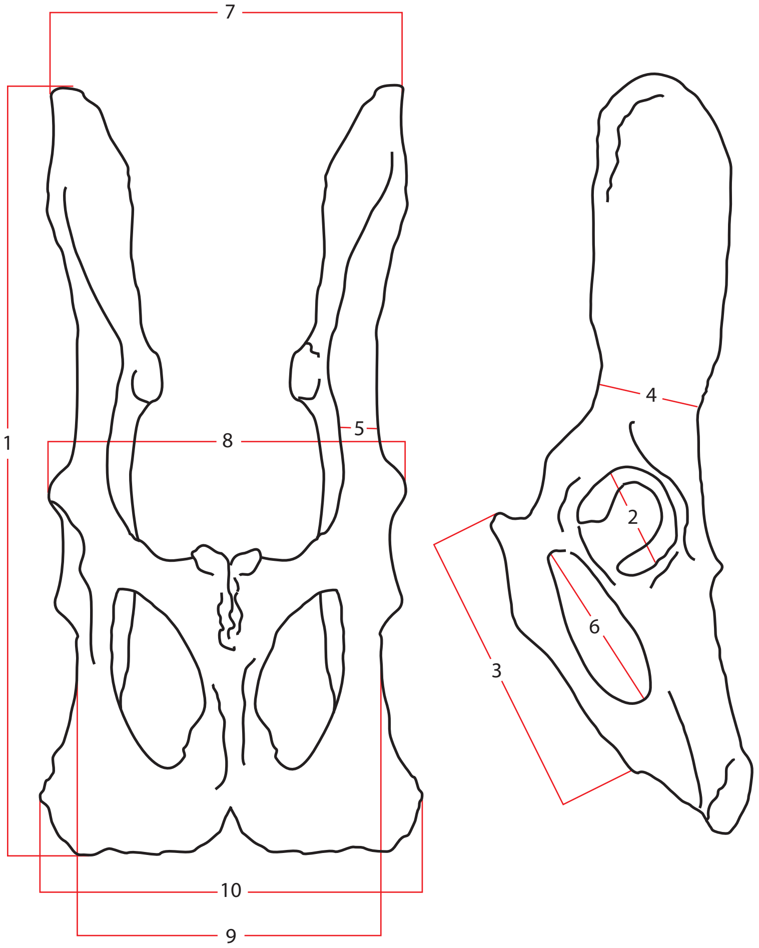

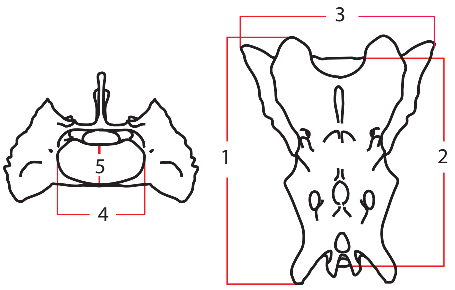

**SACRUM**
1 Gratest length on the ventral side
2 Physiological length
3 Greatest breadth across the wings
4 Greatest breadth of the cranial articular surface
5 Greatest height of the cranial articular surface

**PELVIS**
1 Greatest length of the half
2 Length of the acetabulum on the rim
3 Length of the symphysis (if fused)
4 Smallest height of the shaft of ilium
5 Smallest breadth of the shaft of ilium
6 Inner length of the foramen obturatum
7 Greatest breadth across the iliac crests
8 Greatest breadth across the acetabula
9 Greatest breadth across the ischiatic tuberosity
10 Smallest breadth across the bodies of the ischia

**FORELIMB LONG BONES (Plate 3)**

| HUMERUS | | | |
|---|---|---|---|
| **code** | **description** | **dex** | **sin** |
| 1 | Total length | 185.0 | 185.0 |
| 2 | Anteroposterior breadth of the proximal epiphysis | 40.3 | 38.7 |
| 3 | Transversal breadth of the proximal epiphysis | 30.7 | 30.5 |
| 4 | Transversal breadth of caput (head) | 25.2 | 25.5 |
| 5 | Transversal breadth of the diaphysis (50% length) | 12.5 | 12.6 |
| 6 | Transversal breadth of the distal epiphysis | 37.0 | 37.4 |
| 7 | Greatest breadth of the trochlea | 30.2 | 30.2 |
| 8 | Maximum height of the trochlea | 19.3 | 19.2 |

| ULNA | | | |
|---|---|---|---|
| **code** | **description** | **dex** | **sin** |
| 1 | Total length | 210.5 | 210.5 |
| 2 | Height of the sigmoid cavity | 14.4 | 14.5 |
| 3 | Anteroposterior breadth | 20.7 | 20.7 |
| 4 | Anteroposterior breadth of the sigmoid cavity | 14.3 | 14.4 |
| 5 | Transversal breadth of the olecranon | 14.4 | 13.4 |
| 6 | Anteroposterior breadth of the diaphysis (50% length) | 15.7 | 15.8 |
| 7 | Transversal breadth of the sigmoid cavity | 10.2 | 10.2 |
| 8 | Height of the anterior process of the olecranon | 14.1 | 14.1 |
| 9 | Maximum transversal breadth of the sigmoid cavity | 19.1 | 19.3 |
| 10 | Anteroposterior breadth of the estiloid apophysis | 15.8 | 15.8 |

| RADIUS | | | |
|---|---|---|---|
| **code** | **description** | **dex** | **sin** |
| 1 | Total length | 176.0 | 177.5 |
| 2 | Anteroposterior breadth of the head | 16.6 | 16.6 |
| 3 | Anteroposterior breadth of the neck | 10.3 | 10.3 |
| 4 | Transversal breadth of the head | 11.6 | 11.6 |
| 5 | Anteroposterior breadth of the diaphysis (50% length) | 12.8 | 12.9 |
| 6 | Anteroposterior breadth of the distal epiphysis | 27.3 | 27.1 |
| 7 | Anteroposterior breadth of the distal articular surface | 17.3 | 17.3 |
| 8 | Transversal breadth of the distal epiphysis | 18.3 | 18.3 |
| 9 | Transversal breadth of the distal articular surface | 9.7 | 9.3 |

#### Plate 3
#### FORELIMB LONG BONES

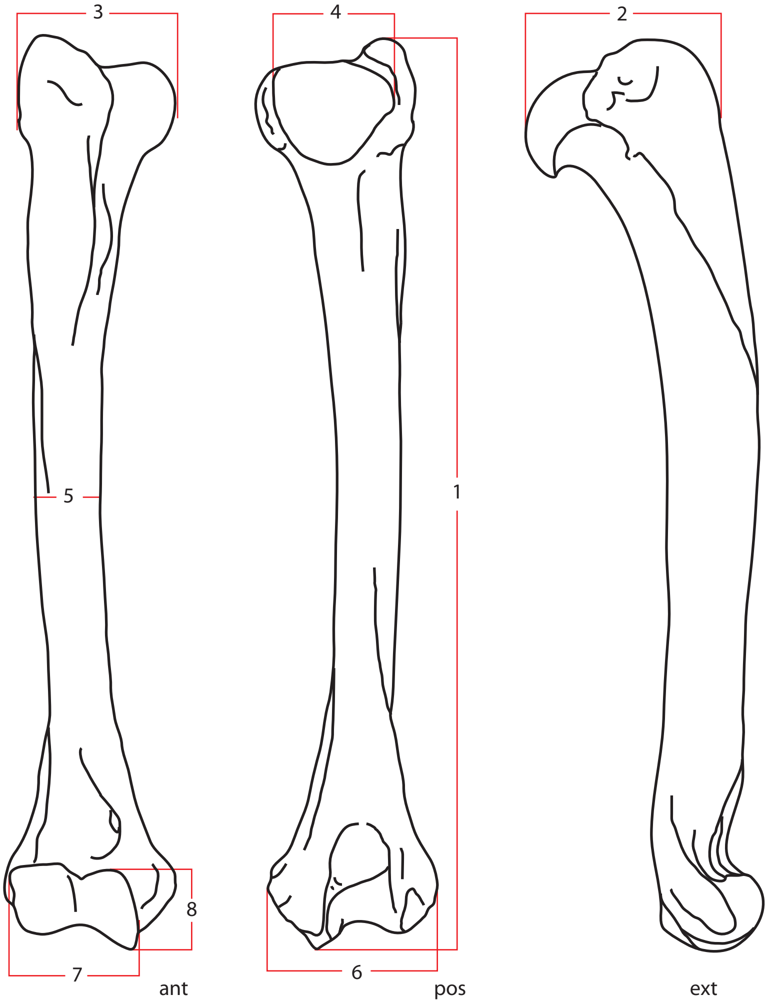

**HUMERUS**
1 Total length
2 Anteroposterior breadth of the proximal epiphysis
3 Transversal breadth of the proximal epiphysis
4 Transversal breadth of caput (head)
5 Transversal breadth of the diaphysis (50% length)
6 Transversal breadth of the distal epiphysis
7 Greatest breadth of the trochlea
8 Maximum height of the throclea

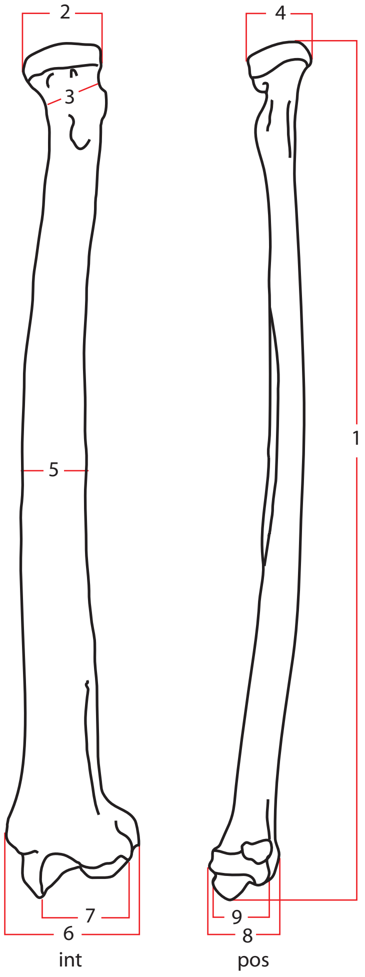

**RADIUS**
1 Total length
2 Anteroposterior breadth of the head
3 Anteroposterior breadth of the neck
4 Transversal breadth of the head
5 Anteroposterior breadth of the diaphysis (50% length)
6 Anteroposterior breadth of the distal epiphysis
7 Anteroposterior breadth of the distal articular surface
8 Transversal breadth of the distal epiphysis
9 Transversal breadth of the distal articular surface

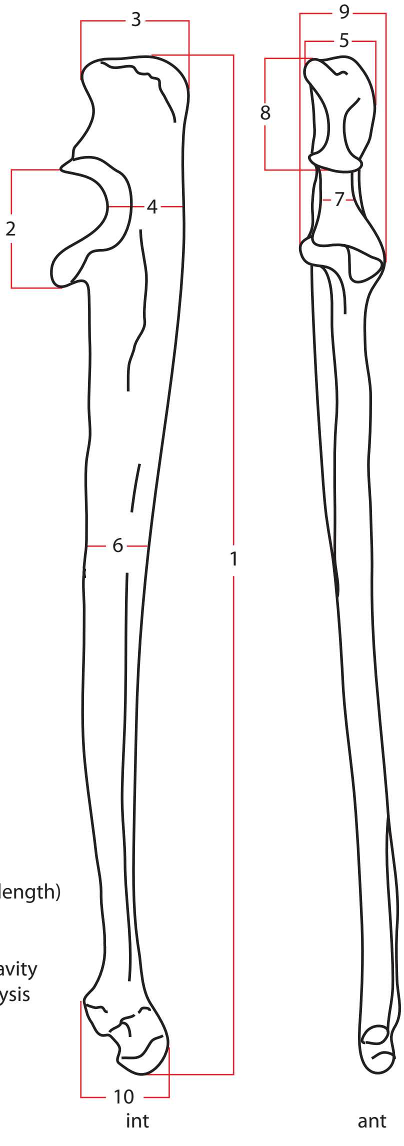

**ULNA**
1 Total length
2 Height of the sigmoid cavity
3 Anteroposterior breadth of the olecranon
4 Anteroposterior breadth of the sigmoid cavity
5 Transversal breadth of the olecranon
6 Anteroposterior breadth of the diaphysis (50% length)
7 Transversal breadth of the sigmoid cavity
8 Height of the anterior process of the olecranon
9 Maximum transversal breadth of the sigmoid cavity
10 Anteroposterior breadth of the estiloid apophysis

**CARPUS, METACARPUS and ANTERIOR PHALANGES (Plate 4)**

| CARPUS | | | |
|---|---|---|---|
| **code** | **description** | **dex** | **sin** |
| **SCAPHOLUNATE** | | | |
| 1 | Transversal diameter | 18.2 | 18.6 |
| 2 | Anteroposterior diameter | 14.6 | 14.6 |
| 3 | Vertical diameter | 9.7 | 9.6 |
| **TRIQUETUM** | | | |
| 1 | Transversal diameter | 12.4 | 12.4 |
| 2 | Anteroposterior diameter | 8.3 | 8.4 |
| 3 | Vertical diameter | 6.3 | 6.3 |
| **PISIFORM** | | | |
| 1 | Transversal diameter | 9.7 | 9.7 |
| 2 | Anteroposterior diameter | 17.9 | 18.0 |
| 3 | Vertical diameter | 9.7 | 9.8 |
| **TRAPEZIUM** | | | |
| 1 | Transversal diameter | 12.3 | 12.4 |
| 2 | Anteroposterior diameter | 7.4 | 7.6 |
| 3 | Vertical diameter | 7.6 | 7.7 |
| **TRAPEZOID** | | | |
| 1 | Transversal diameter | 9.5 | 9.1 |
| 2 | Anteroposterior diameter | 10.3 | 10.1 |
| 3 | Vertical diameter | 5.6 | 5.6 |
| **CAPITATE** | | | |
| 1 | Transversal diameter | 8.1 | 8.3 |
| 2 | Anteroposterior diameter | 13.9 | 14.7 |
| 3 | Vertical diameter | 10.0 | 10.0 |
| **HAMATE** | | | |
| 1 | Transversal diameter | 12.9 | 13.1 |
| 2 | Anteroposterior diameter | 12.6 | 12.4 |
| 3 | Vertical diameter | 9.0 | 8.9 |

| **METACARPALS** | | **I** | | **II** | | **III** | | **IV** | | **V** | |
|---|---|---|---|---|---|---|---|---|---|---|---|
| **code** | **side** | dex | sin | dex | sin | dex | sin | dex | sin | dex | sin |
| 1 | Total length | 20.2 | 20.2 | 60.9 | 60.6 | 70.0 | 69.2 | 65.7 | 65.3 | 53.1 | 53.6 |
| 2 | Proximal transversal breadth | 10.3 | 10.4 | 8.6 | 8.3 | 10.6 | 10.6 | 8.7 | 8.6 | 10.7 | 9.8 |
| 3 | Proximal anteroposterior breadth | 7.7 | 7.6 | 13.1 | 13.2 | 11.3 | 11.6 | 10.2 | 10.8 | 10.3 | 10.4 |
| 4 | Diaphysis transversal breadth | 7.9 | 8.2 | 13.1 | 13.2 | 11.3 | 11.6 | 10.2 | 10.8 | 10.3 | 10.4 |
| 5 | Diaphysis anteroposterior breadth | 5.9 | 6.0 | 6.8 | 7.3 | 6.1 | 6.0 | 6.1 | 5.9 | 5.3 | 5.1 |
| 6 | Distal transversal breadth | 8.6 | 8.6 | 11.2 | 11.3 | 11.7 | 11.8 | 10.7 | 10.7 | 10.8 | 10.3 |
| 7 | Distal anteroposterior breadth | 8.4 | 9.3 | 8.9 | 9.8 | 9.9 | 9.9 | 9.2 | 9.2 | 9.4 | 9.4 |

| **ANTERIOR PHALANGES** | | | | | | | | | | | |
|---|---|---|---|---|---|---|---|---|---|---|---|
| **first phalanx manus** | | **digit I** | | **digit II** | | **digit III** | | **digit IV** | | **digit V** | |
| **code** | **side** | dex | sin | dex | sin | dex | sin | dex | sin | dex | sin |
| 1 | Total length | 15.4 | 15.6 | 29.3 | 29.4 | 35.3 | 35.4 | 33.3 | 33.2 | 25.9 | 25.4 |
| 2 | Proximal transversal breadth | 10.1 | 9.5 | 10.0 | 10.0 | 10.5 | 10.7 | 10.5 | 10.1 | 9.9 | 9.6 |
| 3 | Proximal anteroposterior breadth | 9.5 | 8.3 | 8.6 | 8.7 | 8.3 | 8.0 | 7.7 | 7.7 | 9.5 | 7.7 |
| 4 | Diaphysis transversal breadth | 7.7 | 7.8 | 6.4 | 6.4 | 6,5 | 6,5 | 6.2 | 6.4 | 5.6 | 5.4 |
| 5 | Diaphysis anteroposterior breadth | 5.5 | 5.7 | 5.5 | 5.5 | 4.9 | 4.9 | 5.0 | 5.0 | 5.9 | 5.8 |
| 6 | Distal transversal breadth | 8.2 | 8.3 | 8.1 | 8.1 | 8.4 | 8.5 | 8.2 | 8.2 | 7.6 | 7.5 |
| 7 | Distal anteroposterior breadth | 6.1 | 6.1 | 6.5 | 6.5 | 7.3 | 7.3 | 6.8 | 6.7 | 6.0 | 5.8 |
| **second phalanx manus** | | | | **digit II** | | **digit III** | | **digit IV** | | **digit V** | |
| **code** | side | | | dex | sin | dex | sin | dex | sin | dex | sin |
| 1 | Total length | | | 18.3 | 18.2 | 23.9 | 24.1 | *22,3* | 25,4 | 20,2 | 20,4 |
| 2 | Proximal transversal breadth | | | 8.1 | 8.3 | 8.8 | 8.8 | *9,6* | 8,9 | 8,5 | 8,5 |
| 3 | Proximal anteroposterior breadth | | | 7.1 | 7.2 | 7.5 | 7.6 | *9,0* | 8,2 | 8,6 | 8,5 |
| 4 | Diaphysis transversal breadth | | | 6.1 | 6.3 | 4.8 | 5.0 | *6,5* | 4,5 | 5,8 | 5,1 |
| 5 | Diaphysis anteroposterior breadth | | | 4.7 | 4.7 | 4.8 | 4.8 | *7,0* | 5,3 | 5,6 | 5,7 |
| 6 | Distal transversal breadth | | | 7.0 | 6.8 | 7.1 | 7.1 | 7,6 | 7,6 | 7,9 | 7,9 |
| 7 | Distal anteroposterior breadth | | | 5.9 | 5.6 | 5.8 | 5.6 | 5,8 | 5,9 | 6,3 | 5,9 |
| **third phalanx manus** | | **digit I** | | **digit II** | | **digit III** | | **digit IV** | | **digit V** | |
| **code** | side | dex | sin | dex | sin | dex | sin | dex | sin | dex | sin |
| 1 | Total length * | 19,8 | 18,3 | 18,4 | 18,4 | 22,2 | 22,7 | 21,4 | 20,8 | 19,7 | 20.1 |
| 2 | Proximal transversal breadth | 6.0 | 6.0 | 6,2 | 6,2 | 7,3 | 7,6 | 6,5 | 7,1 | 5,9 | 6,1 |
| 3 | Proximal height | 8,7 | 8,6 | 9 | 8,1 | 11,1 | 11,4 | 9,2 | 10,1 | 8 | 9.2 |

### Plate 4
### CARPUS, TARSUS and PHALANGES

**CARPALS**
1 Transversal diameter
2 Anteroposterior diameter
3 Vertical diameter

**SCAPHOLUNATE**

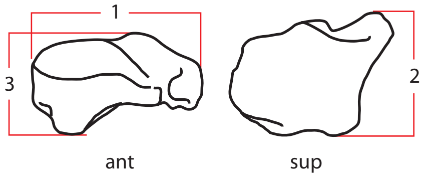

**TRIQUETRUM**

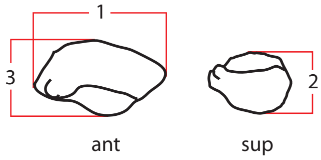

**PISIFORM**

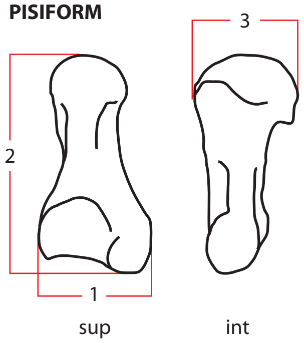

**TRAPEZIUM**

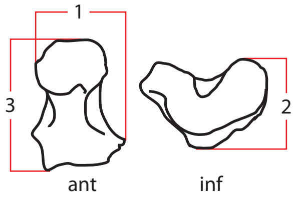

**TRAPEZOID**

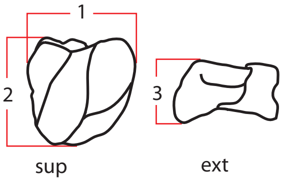

**CAPITATE**

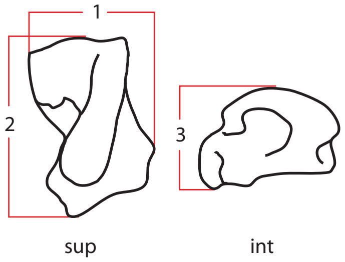

**HAMATE**

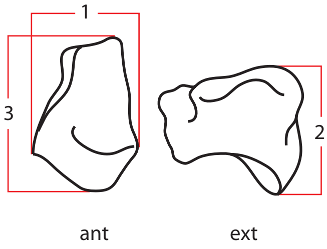

**FIRST METACARPAL, I**

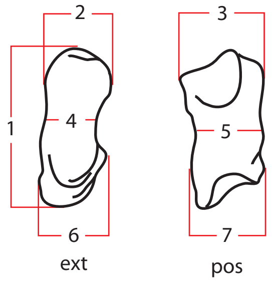

**THIRD METACARPAL, III**

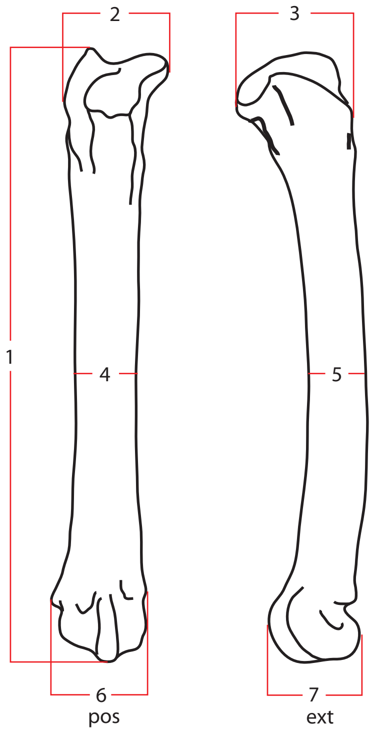

**METACARPALS**
1 Total length
2 Proximal transversal diameter
3 Proximal anteroposterior diameter
4 Diaphysis transversal diameter
5 Diaphysis anteroposterior diameter
6 Distal transversal diameter
7 Distal anteroposterior diameter

**PHALANGES 1, 2**
1 Total length
2 Proximal transversal diameter
3 Proximal anteroposterior diameter
4 Diaphysis transversal diameter
5 Diaphysis anteroposterior diameter
6 Distal transversal diameter
7 Distal anteroposterior diameter

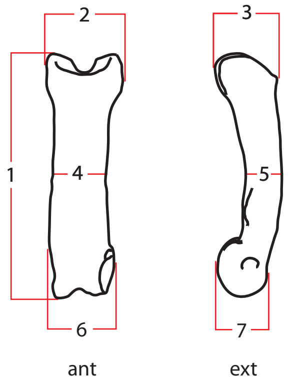

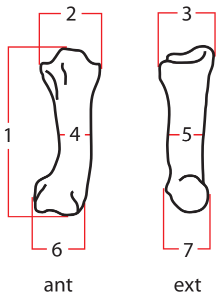

**PHALANGE 3**
1 Total length
2 Proximal transversal diameter
3 Proximal anteroposterior diameter

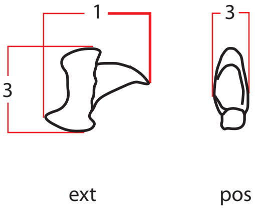

| HINDLIMB LONG BONES & PATELLA (Plate 5) |
|---|

| FEMUR | | | |
|---|---|---|---|
| **code** | **description** | **dex** | **sin** |
| 1 | Total length | 216.0 | 216.0 |
| 2 | Distance between trochleas | 32.7 | 32.7 |
| 3 | Transversal breadth of the proximal epiphysis | 38.9 | 40.0 |
| 4 | Transversal breadth of caput | 17.6 | 17.8 |
| 5 | Transversal breadth of the distal diaphysis | 36.5 | 36.6 |
| 6 | Transversal breadth of the diaphysis (50% length) | 15.0 | 15.0 |
| 7 | Inferior breadth of the distal articular surface | 16.7 | 16.6 |
| 8 | Anteroposterior breadth of the distal epiphysis | 31.3 | 31.4 |
| 9 | Distance between epicondyles | 9.3 | 9.4 |

| PATELLA | | | |
|---|---|---|---|
| **code** | **description** | **dex** | **sin** |
| 1 | Vertical diameter | 28.3 | 28.3 |
| 2 | Transversal diameter | 17.5 | 17.8 |
| 3 | Anteroposterior diameter | 10.8 | 10.8 |

| TIBIA | | | |
|---|---|---|---|
| **code** | **description** | **dex** | **sin** |
| 1 | Total length | 221.5 | 222.0 |
| 2 | Transversal breadth of the proximal epiphysis | 38.0 | 37.5 |
| 3 | Anteroposterior breadth of the proximal epiphysis | 41.7 | 41.1 |
| 4 | Transversal breadth of the diaphysis (50% length) | 13.7 | 13.7 |
| 5 | Anteroposterior breadth of the distal epiphysis | 28.3 | 28.3 |
| 6 | Transversal breadth of the distal epiphysis | 15.7 | 15.2 |

| FIBULA | | | |
|---|---|---|---|
| **code** | **description** | **dex** | **sin** |
| 1 | Total length | 207.5 | 207.0 |
| 2 | Transversal breadth of the proximal epiphysis | 15.6 | 15.4 |
| 3 | Transversal breadth of the distal epiphysis | 15.5 | 15.5 |
| 4 | Transversal breadth of the diaphysis (50% length) | 4.4 | 4.3 |

### Plate 5
### HINDLIMB LONG BONES

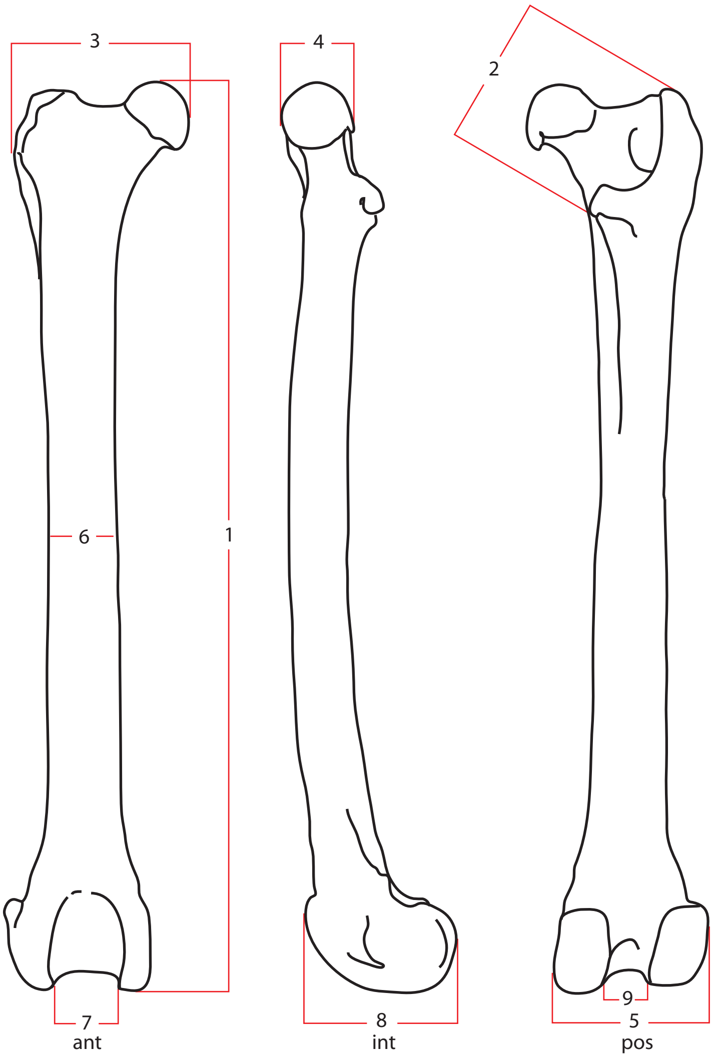

**FEMUR**
1 Total length
2 Distance between trochleas
3 Transversal breadth of the proximal epiphysis
4 Transversal breadth of caput
5 Transversal breadth of the distal diaphysis
6 Transversal breadth of the diaphysis (50% length)
7 Inferior breadth of the distal articular surface
8 Anteroposterior breadth of the distal epiphysis
9 Distance between epicondyles

**PATELLA**
1 Vertical diameter
2 Transversal diameter
3 Anteroposterior diameter

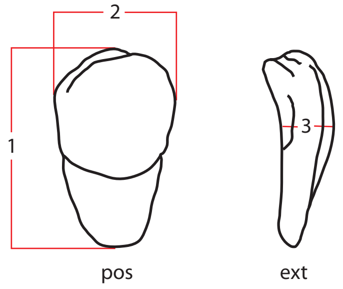

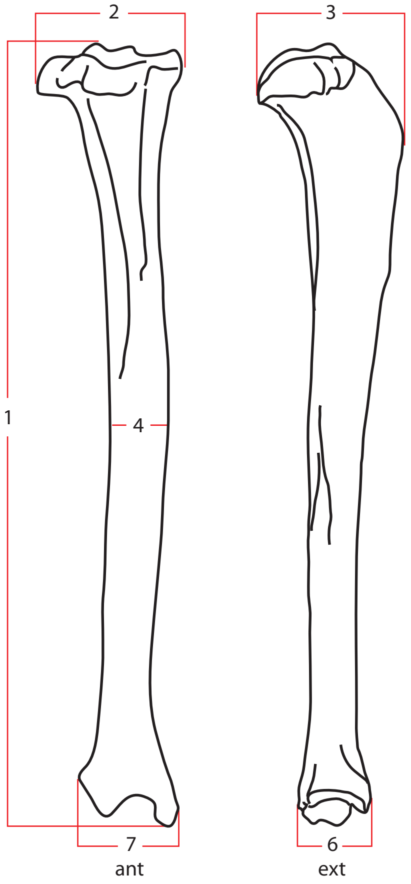

**TIBIA**
1 Total length
2 Transversal breadth of the proximal epiphysis
3 Anteroposterior breadth of the proximal epiphysis
4 Transversal breadth of the diaphysis (50% length)
5 Anteroposterior breadth of the distal epiphysis
6 Transversal breadth of the distal epiphysis

**FIBULA**
1 Total length
2 Transversal breadth of the proximal epiphysis
3 Transversal breadth of the distal epiphysis
4 Transversal breadth of the diaphysis (50% length)

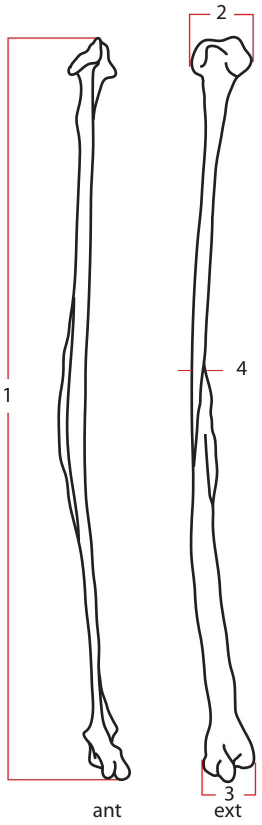

**TARSUS, METATARSUS (Plate 6) and POSTERIOR PHALANGES (as in Plate 4)**

| TARSUS | | | |
|---|---|---|---|
| **code** | **description** | **dex** | **sin** |
| **CALCANEUS** | | | |
| 1 | Transversal diameter | 37.8 | 37.4 |
| 2 | Anteroposterior diameter | 23.5 | 22.9 |
| 3 | Vertical diameter | 56.4 | 56.8 |
| 4 | Transversal diameter of the tuber | 14.5 | 14.6 |
| 5 | Anteroposterior diameter of the tuber | 14.4 | 14.4 |
| **ASTRAGALUS** | | | |
| 1 | Transversal diameter | 22.6 | 22.7 |
| 2 | Anteroposterior diameter | 29.6 | 29.7 |
| 3 | Vertical diameter | 17.8 | 17.9 |
| **CUBOID** | | | |
| 1 | Transversal diameter | 15.0 | 14.8 |
| 2 | Anteroposterior diameter | 14.3 | 13.7 |
| 3 | Vertical diameter | 16.8 | 16.5 |
| **SCAPHOID** | | | |
| 1 | Transversal diameter | 14.1 | 14.1 |
| 2 | Anteroposterior diameter | 19.2 | 19.1 |
| 3 | Vertical diameter | 16.2 | 16.2 |
| **CUNEIFORM 1** | | | |
| 1 | Transversal diameter | 5.9 | 6.1 |
| 2 | Anteroposterior diameter | 8.5 | 8.7 |
| 3 | Vertical diameter | 9.8 | 9.7 |
| **CUNEIFORM 2** | | | |
| 1 | Transversal diameter | 9.3 | 9.5 |
| 2 | Anteroposterior diameter | 5.6 | 5.8 |
| 3 | Vertical diameter | 7.9 | 7.5 |
| **CUNEIFORM 3** | | | |
| 1 | Transversal diameter | 11.6 | 11.6 |
| 2 | Anteroposterior diameter | 20.3 | 20.3 |
| 3 | Vertical diameter | 12.2 | 12.3 |

| **METATARSALS** | | **I** | | **II** | | **III** | | **IV** | | **V** | |
|---|---|---|---|---|---|---|---|---|---|---|---|
| **code** | **side** | dex | sin | dex | sin | dex | sin | dex | sin | dex | sin |
| 1 | Total length | 13.1 | 13.1 | 80.8 | 81.1 | 90.8 | 91.0 | 91.4 | 91.2 | 85.2 | 82.5 |
| 2 | Proximal transversal breadth | 6.2 | 6.1 | 8.6 | 8.5 | 12.9 | 13.0 | 9.1 | 9.2 | 10.3 | 10.4 |
| 3 | Proximal anteroposterior breadth | 5.6 | 5.7 | 12.7 | 13.1 | 16.4 | 16.6 | 13.4 | 11.6 | 8.4 | 8.1 |
| 4 | Diaphysis transversal breadth | | | 7.0 | 6.8 | 9.2 | 9.3 | 6.9 | 7.1 | 5.5 | 5.5 |
| 5 | Diaphysis anteroposterior breadth | | | 6.1 | 6.3 | 6.8 | 6.9 | 7.3 | 7.0 | 4.7 | 4.6 |
| 6 | Distal transversal breadth | | | 12.3 | 12.3 | 12.8 | 13.0 | 11.3 | 11.8 | 10.4 | 10.2 |
| 7 | Distal anteroposterior breadth | | | 10.7 | 10.4 | 11.3 | 11.3 | 10.3 | 10.6 | 9.7 | 9.5 |

| **POSTERIOR PHALANGES** | | | | | | | | | | | |
|---|---|---|---|---|---|---|---|---|---|---|---|
| **first phalanx pes** | | **digit I** | | **digit II** | | **digit III** | | **digit IV** | | **digit V** | |
| **code** | **side** | dex | sin | dex | sin | dex | sin | dex | sin | dex | sin |
| 1 | Total length | 10.9 | 11.0 | 30.7 | 30.7 | 36.7 | 36.7 | 35.7 | 35.5 | 27.5 | 27.6 |
| 2 | Proximal transversal breadth | | | 10.7 | 10.6 | 12.6 | 12.3 | 10.9 | 11.1 | 9.1 | 8.8 |
| 3 | Proximal anteroposterior breadth | | | 9.7 | 9.9 | 9.8 | 9.8 | 9.5 | 9.1 | 8.7 | 8.6 |
| 4 | Diaphysis transversal breadth | | | 6.5 | 6.6 | 7.9 | 7.7 | 7.2 | 7.9 | 5.3 | 5.1 |
| 5 | Diaphysis anteroposterior breadth | | | 6.8 | 6.8 | 5.5 | 5.8 | 5.7 | 5.8 | 5.7 | 5.8 |
| 6 | Distal transversal breadth | | | 9.1 | 8.8 | 9.7 | 9.5 | 9.0 | 9.1 | 7.6 | 7.8 |
| 7 | Distal anteroposterior breadth | | | 6.6 | 6.6 | 7.6 | 7.7 | 7.3 | 7.2 | 5.9 | 5.9 |
| **second phalanx pes** | | | | **digit II** | | **digit III** | | **digit IV** | | **digit V** | |
| **code** | **side** | | | dex | sin | dex | sin | dex | sin | dex | dex |
| 1 | Total length | | | 18,4 | 19,5 | 26,0 | 26,0 | 25,2 | 24,8 | 18,8 | 20,9 |
| 2 | Proximal transversal breadth | | | 8,1 | 9,3 | 9,9 | 10,1 | 9,4 | 9,4 | 8,1 | 9,2 |
| 3 | Proximal anteroposterior breadth | | | 8,0 | 8,3 | 8,5 | 8,5 | 7,9 | 8,1 | 7,0 | 8,1 |
| 4 | Diaphysis transversal breadth | | | 5,5 | 6,4 | 5,5 | 6,4 | 6,2 | 6,1 | 5,7 | 5,8 |
| 5 | Diaphysis anteroposterior breadth | | | 4,4 | 6,1 | 5,9 | 6,0 | 5,3 | 5,2 | 4,8 | 5,7 |
| 6 | Distal transversal breadth | | | 7,0 | 8,0 | 8,1 | 8,0 | 7,6 | 7,6 | 6,8 | 8,1 |
| 7 | Distal anteroposterior breadth | | | 5,7 | 6,0 | 6,1 | 6,1 | 6,2 | 6,3 | 5,6 | 6,2 |
| **third phalanx pes** | | | | **digit II** | | **digit III** | | **digit IV** | | **digit V** | |
| **code** | **side** | | | dex | sin | dex | sin | dex | sin | dex | sin |
| 1 | Total length * | | | 18.3 | 18.1 | 19.2 | 18.9 | 18.0 | 16.8 | 15.9 | 15.8 |
| 2 | Proximal transversal breadth | | | 6.5 | 6.5 | 6.9 | 6.9 | 6.7 | 6.8 | 6.1 | 5.8 |
| 3 | Proximal height | | | 8.8 | 8.5 | 9.0 | 8.8 | 8.4 | 8.6 | 7.5 | 7.7 |

### Plate 6
### TARSUS, METATARSUS

**CALCANEUS, CUBOID, NAVICULAR, CUNEIFORMS**
1 Transversal diameter
2 Anteroposterior diameter
3 Vertical diameter

**TALUS**
1 Transversal diameter
2 Anteroposterior diameter
3 Vertical diameter
4 Tuber transversal diameter
5 Tuber anteroposterior diameter

**CALCANEUS**

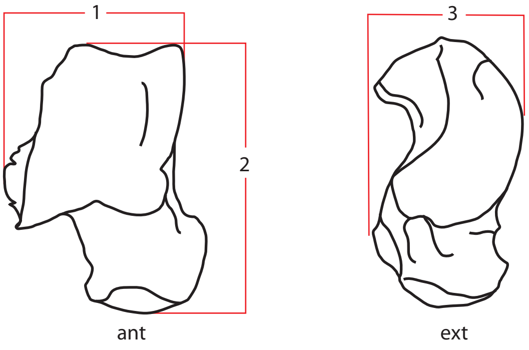

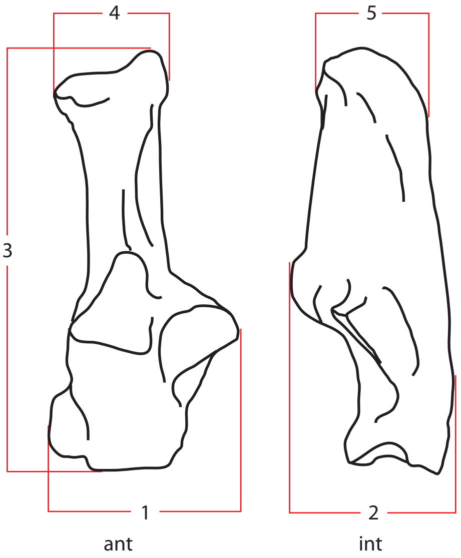

**CUBOID**

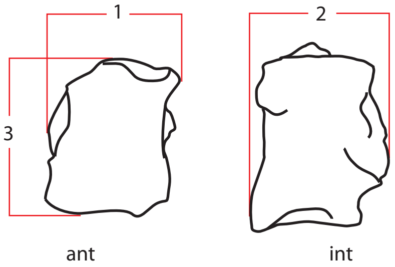

**NAVICULAR**

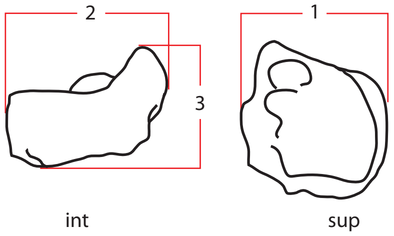

**INTERNAL CUNEIFORM**

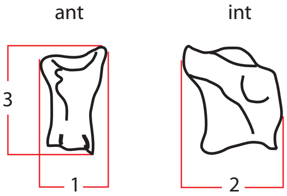

**MEDIAL CUNEIFORM**

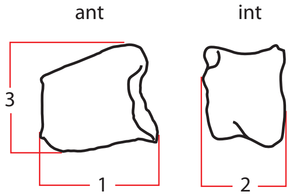

**EXTERNAL CUNEIFORM**

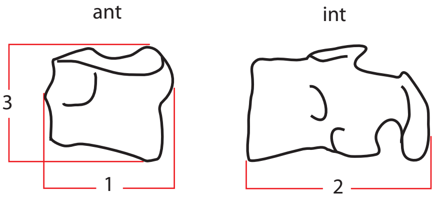

**THIRD METATARSAL, III**

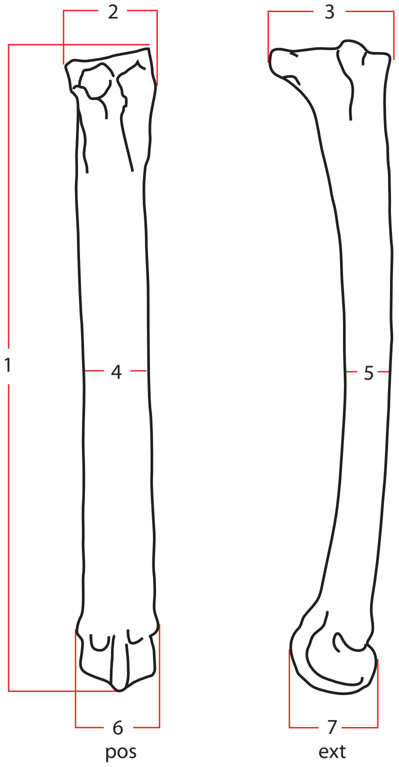

**FIRST METATARSAL, I**

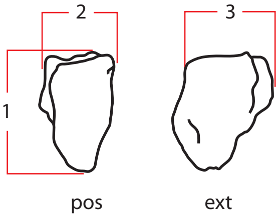

**METATARSALS**
1 Total length
2 Proximal transversal diameter
3 Proximal anteroposterior diameter
4 Diaphysis transversal diameter
5 Diaphysis anteroposterior diameter
6 Distal transversal diameter
7 Distal anteroposterior diameter
